# *CSLB4*-mediated cell wall remodeling decouples phloem access from aphid performance in *Arabidopsis thaliana*

**DOI:** 10.64898/2026.05.06.723330

**Authors:** Felipe Moraga, Daniela Arias-G, Dayan Sanhueza, Joaquín Delgado-Rioseco, Isabel Fuenzalida-Valdivia, Catalina Inostroza-Aguirre, Micaela Peppino-Margutti, Matias Ramos, Diego Zavala-Torres, Francisca Ormeño, Romina V. Sepúlveda, Corina Fusari, Ariel Herrera-Vásquez, Susana Saez-Aguayo, Francisca Blanco-Herrera

**Affiliations:** Centro de Biotecnología Vegetal, Facultad de Ciencias de la Vida, Universidad Andrés Bello, Santiago, Chile; Centro de Mejoramiento Genético y Fenómica Vegetal, Facultad de Ciencias Agrarias, Universidad de Talca, Talca, Chile; Facultad de Ciencias Biológicas, Pontificia Universidad Católica de Chile, Santiago, Chile; Center for Bioinformatics and Integrative Biology (CBIB), Facultad de Ciencias de la Vida, Universidad Andrés Bello, Santiago, Chile; ANID-Millennium Nucleus in Data Science for Plant Resilience (PhytoLearning), Facultad de Ciencias de la Vida, Universidad Andres Bello, Santiago, Chile; Centro de Estudios Fotosintéticos y Bioquímicos (CEFOBI, CONICET/UNR). Facultad de Ciencias Bioquímicas y Farmacéuticas, Universidad Nacional de Rosario (FBioyF, UNR), Rosario, Argentina; Center for Advancing Agri-Food System Transformation (CA2FST), Santiago, Chile; Millennium Science Initiative Program (ANID), Millennium Institute for Integrative Biology (iBio), Santiago, Chile

**Keywords:** *Arabidopsis thaliana*, *Brevicoryne brassicae*, cell wall integrity, cellulose synthase-like B (CSLB), genome-wide association study (GWAS), haplotype analysis, phloem feeding, plant–insect interaction

## Abstract

The plant cell wall (CW) is a key determinant of plant defense; however, the extent to which natural variation in CW architecture contributes to resistance against phloem-feeding insects remains unclear. Here, we combined genome-wide association studies (GWAS) with functional analyses to identify genetic determinants of resistance against the specialist aphid *Brevicoryne brassicae* in *Arabidopsis thaliana*. GWAS conducted on 200 natural accessions identified a single locus on chromosome 2 associated with aphid performance. Integration of haplotype and epidermis-specific expression data prioritized *CSLB4*, a member of the cellulose synthase-like B family. Loss-of-function *cslb4* mutants showed reduced aphid offspring, indicating enhanced resistance to *B. brassicae*, whereas performance of the generalist aphid *Myzus persicae* was unaffected. Electrical penetration graph analyses revealed earlier phloem access on *cslb4* mutants despite reduced performance, indicating a decoupling between phloem access and aphid success. Biochemical and immunolocalization analyses showed that *CSLB4* disruption altered CW architecture, including increased xyloglucan epitope accessibility in mesophyll cell walls and reduced callose deposition upon aphid infestation. In addition, CSLB4 localized to Golgi-associated compartments, and *in silico* analyses are consistent with a role in non-cellulosic polysaccharide biosynthesis. Together, these findings identified *CSLB4* as a modulator of CW architecture that uncouples phloem access from aphid performance.

**Highlight:** A GWAS identifies *CSLB4* as a regulator of cell wall architecture that uncouples aphid-feeding from performance, revealing a new mechanism of plant resistance to specialist insects.

## Introduction

Plant cells are surrounded by the cell wall (CW), a dynamic extracellular matrix that functions not only as a structural barrier but also as a key regulator of plant immunity to biotic stress (Somerville *et al*., 2004; Bacete *et al*., 2018). The CW also acts as a reservoir of signaling molecules and antimicrobial compounds that contribute to the activation of immune responses (Miedes *et al*., 2014; Molina *et al*., 2024b). Perturbation in CW architecture can compromise cell wall integrity (CWI), activating surveillance mechanisms that detect structural changes and trigger defense signaling (Bacete and Hamann, 2020; Wan *et al*., 2021). These include the perception of damage-associated molecular patterns (DAMPs), leading to immune signaling pathways that overlap with classical pathogen-triggered immunity (Hahn *et al*., 1981; Yang *et al*., 2021; Molina *et al*., 2024a).

Increasing evidence indicates that CW remodeling plays a critical role in pathogen colonization success. During pathogen attack, pathogens actively modify the CW through the secretion of effectors, including cell wall–degrading enzymes (CWDEs), to facilitate tissue penetration and nutrient acquisition (Silva-Sanzana *et al*., 2020; Bradley *et al*., 2022). Notably, successful pathogens must finely regulate this activity to avoid excessive release of immunogenic fragments (Yang *et al*., 2021; Rebaque *et al*., 2025), reflecting a co-evolutionary interplay between plants and pathogens.

At the same time, plants dynamically regulate CW architecture through endogenous cell wall–modifying enzymes, reinforcing or altering CW properties as part of their defense strategy during plant–pathogen interactions (Bethke *et al*., 2016; Silva-Sanzana *et al*., 2019; Mbiza *et al*., 2022). In particular, pectin biosynthesis has emerged as a key contributor to CWI and immune activation (Bethke *et al*., 2016), while modulation of pectin methylesterification influences plant defenses against phloem-feeding insects such as *Myzus persicae* (Silva-Sanzana *et al*., 2019).

The outcome of these interactions depends on the balance between CW degradation and reinforcement, ultimately determining plant resistance or susceptibility. Changes in CW composition can affect both mechanical properties and CWI-mediated signaling (Houston *et al*., 2016; Bacete and Hamann, 2020), thereby influencing insect feeding behavior and plant susceptibility. However, the genetic and structural determinants linking CW architecture to aphid resistance remain poorly understood. We hypothesized that CW architecture independently modulates early probing events and downstream determinants of aphid performance.

Hemicelluloses are key components of the primary CW that interact with cellulose microfibrils, contributing to CW architecture by tethering microfibrils and modulating mechanical properties. These interactions strongly influence the physical properties of the CW and its ability to respond to external stress (Hamann, 2015; Vaahtera *et al*., 2019). Hemicellulose biosynthesis is mediated by members of the Cellulose Synthase-Like (CSL) gene family (Richmond and Somerville, 2000; Lerouxel *et al*., 2006), which has undergone extensive diversification in plants (Yin *et al*., 2009; Little *et al*., 2018). While several CSL subfamilies have been functionally characterized (Liepman *et al*., 2005; Burton *et al*., 2006; Cocuron *et al*., 2007; Doblin *et al*., 2009; Yang *et al*., 2020), the biological roles of others, including the CSLB family, remain largely unexplored, particularly in the context of plant–insect interactions.

Functional studies have shown that different CSL subfamilies synthesize distinct hemicellulosic polysaccharides, including mannans and glucomannans (CSLA) (Dhugga *et al*., 2004; Liepman *et al*., 2005), xyloglucan backbones (CSLC) (Cocuron *et al*., 2007; Dwivany *et al*., 2009), cellulose-like β-glucans (CSLD) (Yang *et al*., 2020), and mixed-linkage β-glucans in grasses (CSLF and CSLH) (Burton *et al*., 2006; Doblin *et al*., 2009). Beyond their structural roles, CSL-mediated modifications of the CW have been implicated in responses to biotic and abiotic stresses, including osmotic stress (Zhu *et al*., 2010), as well as in mediating trade-offs between plant growth and defense (Liu *et al*., 2022). Emerging evidence also indicates that certain CSL members (e.g., CSLM) contribute to the glycosylation of specialized metabolites, such as triterpenoid saponins, influencing susceptibility to chewing insects (Jozwiak *et al*., 2020, 2024).

Despite these advances, the CSLB family remains poorly characterized, with limited functional evidence beyond sequence-based annotation, predicted enzymatic activity, and phylogenetic placement within the glycosyltransferase family (GT2) (Saxena *et al*., 1995; Yin *et al*., 2014; Little *et al*., 2018). In *Arabidopsis thaliana*, the CSLB family comprises six members (CSLB1–CSLB6), which share moderate to high sequence similarity (>68% at the protein level) and likely arose through gene duplication events. Such duplication is a major driver of gene family expansion in plants, generating paralogous genes that may undergo functional diversification, including partial redundancy or spatial and/or temporal specialization (Lin *et al*., 1999; Panchy *et al*., 2016). However, direct evidence linking CSLB function to CW-mediated interactions with phloem-feeding insects is still lacking.

Natural genetic variation provides a powerful framework for uncovering genes underlying complex traits (Atwell *et al*., 2010; Li *et al*., 2010; Kloth *et al*., 2016), yet its potential to identify genetic determinants of insect resistance remains incomplete. In particular, how CW properties influence herbivory at the genetic level is not well understood. Genome-wide association studies (GWAS) therefore offer an unbiased approach to identifying loci associated with variation in aphid performance across genetically diverse populations, including the cabbage aphid, *B. brassicae*, a major pest of Brassicaceae crops (Blackman and Eastop, 2000; Van Emden and Harrington, 2007). Here, we performed GWAS in *A. thaliana* to identify loci associated with resistance to the specialist aphid *B. brassicae*. We identified a locus containing a previously uncharacterized member of the CSLB family and demonstrated that disruption of *CSLB4* alters aphid performance, feeding behavior, and CW architecture. Loss of *CSLB4* function reduces aphid offspring production and alters feeding dynamics, revealing a decoupling between phloem access and aphid performance. These findings challenge the assumption that efficient phloem access predicts aphid performance and instead highlight the importance of post-access factors, likely associated with CW-mediated properties or phloem quality, in determining herbivore success. Together, our results uncover a novel role for CSLB-mediated CW remodeling in plant–aphid interactions and identify *CSLB4* as a negative regulator of resistance to *B. brassicae*.

## Materials and methods

### Plant materials, insects, and growth conditions

*Arabidopsis thaliana* natural accessions (*n* = 200; **Supplementary Table S1**) and T-DNA insertional lines *cslb4-1* (SALK_067582) and *cslb4-2* (SALK_118481) were obtained from the Arabidopsis Biological Resource Center (ABRC) (Alonso *et al*., 2003). The Columbia-0 (Col-0) accession was used as the wild-type (WT) background for all mutant analyses. Homozygous insertion lines were confirmed by PCR genotyping using gene-specific and T-DNA border primers. Primer sequences used for genotyping are listed in **Supplementary Table S2**. For GWAS phenotyping, seeds were stratified at 5 °C for 5 days and grown in 5-cm pots filled with a 1:1 (v/v) mixture of organic soil (Kekkilä, Finland) and vermiculite (Protekta, Chile) under controlled conditions (12 h light/12 h dark photoperiod, 22 °C, 22 °C, 60% relative humidity). Plants were thinned out to one seedling per pot two weeks after germination and fertilized weekly with a nutritive solution (Phostrogen®, UK) until the experiments were performed. For the *in vitro* aphid preference assay, seeds were surface sterilized with 30% hypochlorite, stratified for 24 h at 5 °C, and sown on half-strength Murashige and Skoog (MS) medium with vitamins supplemented with 1% (w/v) sucrose and 0.8 % (w/v) agar (pH 5.7). Plates were maintained under the same growth conditions described above.

Colonies of the specialist aphid *Brevicoryne brassicae* and the generalist aphid *Myzus persicae* were maintained on kale plants (*Brassica oleracea* var. sabellica) in a climate-controlled greenhouse under a 12 h light/12 h dark photoperiod at 22 °C. Aphid colonies were regularly transferred to fresh plants to maintain vigorous populations and ensure consistent physiological status for all experiments.

### Aphid resistance assays (no-choice)

Aphid reproductive performance (offspring number) was evaluated under no-choice conditions using 5-week-old plants. Two neonate nymphs (founders), obtained from synchronized adult females maintained on kale, were confined to each plant (schematic representation of the experimental design in **Supplementary Fig. S1**). For *B. brassicae*, the total number of offspring was recorded 19 days after infestation, whereas for *M. persicae*, offspring were counted 14 days after infestation to account for differences in life cycle duration between species. Each biological replicate consisted of a single plant infested with two founder nymphs.

For GWAS phenotyping, the 200 natural accessions were arranged in an incomplete block design with four biological replicates per accession, two blocks per replicate, and 100 genotypes per block. Offspring number per plant was used as a quantitative measure of aphid resistance. For functional validation, aphid performance was assessed on Col-0 and *cslb4* mutant lines using 15–20 biological replicates per genotype per experiment. A second independent T-DNA insertion line (*cslb4-2*) was included in no-choice assays to confirm the phenotype associated with *CSLB4* disruption.

### Electrical penetration graph (EPG) analysis

Aphid-feeding behavior was analyzed using the EPG technique on 5-week-old Col-0 and *cslb4-1* plants. Adult apterous *B. brassicae* were immobilized, and a thin gold wire (18 μm diameter, approximately 1.5 cm long) was attached to the dorsum using a water-based silver glue (EPG Systems, The Netherlands). The opposite end of the wire was connected to the probe of a DC Giga-8 EPG system (EPG Systems, The Netherlands), and a copper electrode was inserted into the soil to complete the electrical circuit.

Each wired aphid was placed on an individual plant, and stylet activity was recorded for 8 h, with 24 biological replicates per genotype (each replicate consisted of a single aphid monitored on a single plant). Waveforms were annotated using EPG Stylet+ software and further processed with the NPAC-EPGv Excel workbook (Garzo *et al*., 2024). Standard EPG waveforms, including pathway phase C, potential drops (pd), and phloem-related phases (E1 and E2), were identified following established conventions for aphid EPG analysis (Tjallingii, 1988, 1990), and used to derive parameters describing probing behavior, phloem access, and ingestion phases for statistical analysis. Events interrupted at the end of the recording were included in duration calculations, whereas non-occurring waveforms were treated as missing values for specific parameters.

### Statistical modelling of phenotypic data and heritability estimation

Phenotypic data were analyzed using linear mixed models (LMMs) fitted with the *lme4* and *inti* packages in R (version 4.4.0) (R Core Team, 2024) to estimate genotypic effects. Replication and block within replication were treated as random effects, whereas genotype was modeled as a fixed effect to obtain best linear unbiased estimates (BLUEs), which were used for genome-wide association (GWAS) analysis.

Broad-sense heritability (*H*²) was estimated as described in Cullis *et al*. (2006). Estimates were based on the prediction error variance of genotype best linear unbiased predictors (BLUPs), derived from the same LMM by modeling genotype as a random effect, providing a measure of the genetic contribution to phenotypic variation.

### Genome-wide association mapping

Genome-wide association mapping was performed using an LMM implemented in the Genome-wide Efficient Mixed Model Association (GEMMA) algorithm. Phenotypic BLUEs were used as input for association analysis. Genotype data consisting of approximately 3 million SNP markers were obtained from the imputed SNP dataset of the Arabidopsis regional mapping population (Arouisse *et al*., 2020). SNPs with a minor allele frequency (MAF) lower than 0.01 were filtered out prior to association analysis to remove rare variants. After filtering, the final genotype dataset comprised 2,671,481 SNPs.

The LMM incorporated a centered relatedness (kinship) matrix to account for genetic relatedness among accessions. Significance of SNP–trait associations was assessed using the Wald test. Genome-wide significance was determined using a Bonferroni correction threshold of *p* = 1.87 × 10, based on the number of SNPs retained after filtering. Association mapping was conducted using the *vcf2gwas* pipeline, implemented as a Python command-line tool (Vogt *et al*., 2021). Manhattan plots were generated to visualize genome-wide association signals.

### Haplotype analysis and candidate gene identification

To further investigate genomic regions associated with significant SNPs, haplotype analysis was performed within a 50 kb window centered on each lead SNP (±25 kb upstream and downstream). In addition to genome-wide significant loci, regions showing suggestive association signals (−log_10_(*p*) > 7) were also included in the analysis.

Haplotype blocks were inferred using a sliding window of 10 consecutive SNP markers, independently of local linkage disequilibrium (LD) structure. Association between haplotypes and aphid reproductive performance (offspring number) was assessed using *haplo.stats* package in R (R Core Team, 2024). Phenotypic BLUEs from the LMM were used as quantitative trait values. A generalized linear model framework was used to estimate global score statistics and haplotype-specific effects. Differences among haplotype classes were assessed using one-way ANOVA followed by Tukey’s HSD test for multiple comparisons. To further refine the association signal, LD analysis was performed for the most significant locus by assessing pairwise LD (r^2^, coefficient of correlation) between the lead SNP and neighboring variants using R packages *genetics* for calculation and *LDheatmap* for visualization. Candidate genes within the defined genomic intervals were identified based on the *A. thaliana* reference genome (TAIR10), and functional annotations were retrieved from The Arabidopsis Information Resource (TAIR).

### Cell wall composition analysis (AIR and monosaccharide analysis)

Cell wall material was isolated from whole rosettes of 5-week-old Col-0 and *cslb4-1* plants (n = 3 biological replicates; each replicate consisted of pooled rosettes from six independent plants). The tissue was frozen and ground in liquid nitrogen using a mortar and pestle. Alcohol-insoluble residues (AIR) were prepared by sequential extraction with 80% (v/v) ethanol, followed by two 2-hour incubations in methanol:chloroform (1:1, v/v) to remove lipids, and subsequently two 45-minute washes with acetone. The dried AIR was used for monosaccharide analysis.

For cell wall fractionation, AIR samples were sequentially extracted to obtain pectin- and hemicellulose-enriched fractions. Pectin-enriched fractions were obtained using imidazole buffer and ammonium oxalate extraction, as described in Sanhueza *et al*. (2024b). Pectins were extracted by incubating AIR twice with 0.5 M imidazole (pH 7.0) at room temperature, followed by two additional incubations with 0.2 M ammonium oxalate (pH 4.3) at 60 °C. The resulting supernatants were pooled, dialyzed against water using a 12-kDa molecular weight cut-off membrane, and freeze-dried to obtain the pectin-enriched fraction. The residual pellet was used for hemicellulose extraction. It was first rinsed with water and then incubated overnight at 37 °C with shaking in 6 M NaOH containing 1% (w/v) NaBH□. This extraction was performed twice, and the collected supernatants were pooled, dialyzed, and freeze-dried to obtain the hemicellulose-enriched fraction. These sequential extractions yield operationally defined fractions enriched in pectins and hemicelluloses, respectively.

Samples were hydrolyzed with 400 µL of 2 M trifluoroacetic acid (TFA) at 121 °C for 1 hour (AIR) or 45 minutes (pectin and hemicellulose). After hydrolysis, TFA was evaporated under nitrogen at 42 °C. The residues were washed twice with 400 µL of 100% isopropanol, resuspended in 600 µL of Milli-Q water, sonicated for 15 minutes, centrifuged for 1 minute at maximum speed, and filtered through a 0.22 µm syringe filter prior to analysis by HPAEC-PAD. Each hydrolysate was analyzed in technical replicates to ensure analytical reproducibility. Myo-inositol and allose (250 µM each) were used as internal standards and subjected to the same hydrolysis conditions as the samples. Monosaccharides were quantified using a Dionex ICS-3000 ion chromatography system equipped with a pulsed amperometric detector, a CarboPac PA1 (4 × 250 mm) analytical column, and a CarboPac PA1 (4 × 50 mm) guard column, as described in Sanhueza *et al*. (2024b).

Neutral and acidic monosaccharides were separated by HPAEC-PAD at 30 °C using sequential elution. Neutral sugars were resolved isocratically with 20 mM NaOH (1 mL min□¹, 24 min), followed by separation of acidic sugars using 75 mM sodium acetate in 150 mM NaOH (20 min). After each run, the column was washed with 200 mM NaOH and re-equilibrated with 20 mM NaOH (Saez-Aguayo *et al*., 2021). Quantification was performed using calibration curves generated with monosaccharide standards, including neutral sugars [fucose (Fuc), rhamnose (Rha), arabinose (Ara), galactose (Gal), glucose (Glc), xylose (Xyl), and mannose (Man)] and acidic sugars [galacturonic acid (GalA) and glucuronic acid (GlcA)]. Monosaccharide content was expressed as mg g□¹ AIR. Additionally, molar ratios were calculated to assess changes in cell wall composition. For pectin-enriched fractions, the ratios Rha/GalA, Ara/Rha, and Gal/Rha were determined, while the Xyl/Glc ratio was calculated for hemicellulose-enriched fractions. Ratios derived from pectin-enriched fractions are commonly used as proxies for the composition of the rhamnogalacturonan I (RG-I) side chain.

### Quantification of methanol content

Methanol content was determined in pectin fractions derived from AIR samples described above using a modified protocol based on (Anthon and Barrett, 2004). Pectin solutions were prepared at 3 mg mL□¹, and 50 μL aliquots were treated with 50 μL of 0.2 M NaOH and incubated on ice at 4 °C for 1 hour to allow saponification. The reaction was terminated by adding 50 μL of 0.2 M HCl, and the reaction mixture was adjusted to a final volume of 300 μL with Milli-Q water.

For methanol detection, 50 μL of each saponified sample was mixed with 100 μL of 200 mM Tris-HCl (pH 7.5), 40 μL of 3 mg mL□¹ 3-methyl-2-benzothiazolinone hydrazone (MBTH), and 20 μL of alcohol oxidase from *Pichia pastoris* (0.02 U μL□¹; Sigma A-2404), and incubated at 30 °C for 20 minutes, followed by colorimetric detection with 200 μL of a solution containing sulfamic acid and ammonium ferric sulfate dodecahydrate (0.5% w/v each), followed by incubation at room temperature for 20 minutes. The reaction was then diluted with 600 μL of Milli-Q water, and absorbance was measured at 620 nm.

Methanol content was calculated using a standard curve, and values were used as an indirect estimate of pectin methylesterification. All measurements were performed in five technical replicates to ensure analytical reproducibility.

### Immunodot blot assay

Immunodot blot assays were performed using pectin- and hemicellulose-enriched fractions obtained from AIR samples described above, following the methodology described in Sanhueza *et al*. (2024a). Serial dilutions were prepared, and 0.8 µL of each dilution was spotted onto 0.45 µm nitrocellulose membranes (Thermo Scientific, cat. no. 88018). Membranes were blocked using TBS-T buffer (25 mM Tris, 0.15 M NaCl, and 0.1% w/v Tween-20) supplemented with 3% (w/v) skim milk. Primary antibodies were diluted 1:50 in TBS-T containing 1% (w/v) skim milk and incubated at room temperature with the membranes. The 2F4 antibody was prepared in TCaNa buffer (20 mM Tris-HCl, pH 8.2, 0.5 mM CaCl□, and 150 mM NaCl) according to the manufacturer’s instructions and supplemented with 1% (w/v) skim milk and 0.1% (w/v) TBS-T. After primary incubation, membranes were treated with the corresponding alkaline phosphatase-conjugated secondary antibody diluted 1:2000 in TBS-T. Signal detection was performed using BCIP/NBT 1-Step chromogenic substrate (Thermo Scientific, cat. no. 34042).

Dot intensities were quantified using Fiji (ImageJ, National Institutes of Health, USA). Signal intensity was used as a proxy for the relative abundance of specific cell wall epitopes. Three technical replicates were included for each assay.

### Immunofluorescence analysis of xyloglucan using confocal microscopy

Leaf samples from 5-week-old *A. thaliana* plants (Col-0 and *cslb4-1*) were fixed in FAA solution (10% formaldehyde, 5% acetic acid, and 50% ethanol) for 24 h. Samples were dehydrated through a graded ethanol series (10–100%) at 4 °C under vacuum infiltration and embedded in LR White resin (Sigma-Aldrich, USA) using increasing resin–ethanol mixtures (1:2, 1:1, and 2:1), followed by two incubations in 100% resin, as previously described (Sanhueza *et al*., 2024b), with minor modifications. Polymerization was performed for 24 h at 60 °C. Semi-thin transverse sections (5 µm thickness) were obtained using a rotary microtome and mounted on charged microscope slides for immunolabeling. Sections were incubated with the rat monoclonal antibody LM25 (1:50; Sigma-Aldrich, USA), which recognizes XXXG-, XXLG-, and XLLG-type xyloglucan epitopes (Pedersen *et al*., 2012), followed by incubation with an Alexa Fluor 488-conjugated anti-rat IgG secondary antibody (1:500). Cell walls were counterstained with Calcofluor White (Sigma-Aldrich, USA).

For each biological replicate (n = 3; each replicate consisted of a single plant, from which two leaves were collected), three transverse sections were prepared per leaf, and one section per biological replicate was selected for imaging based on section quality and tissue integrity. Images were acquired using a Leica TCS LSI confocal laser-scanning microscope (Leica Microsystems, Germany) with identical acquisition settings across all samples. Fluorescence signal was quantified across the entire section area. Regions of interest corresponding to mesophyll and epidermal cell walls were manually defined, and mean pixel intensity was determined after background subtraction using Fiji, as described in Silva-Sanzana *et al*. (2019). In addition, the area of antibody-derived signal (green channel) was quantified and expressed as a percentage of the total cell wall area, defined by the Calcofluor White signal (magenta channel). Signal intensity was interpreted as a proxy for the relative abundance or accessibility of xyloglucan epitopes.

### Callose staining and quantification

Callose deposition was analyzed in 5-week-old *A. thaliana* plants (Col-0 and *cslb4-1*) after aphid infestation. For infestation assays, approximately 40 *B. brassicae* aphids were confined to two fully expanded leaves per plant using clip cages for 48 h. Leaves were fixed in acetic acid:ethanol (1:3, v/v) and washed in 150 mM K□HPO□ buffer. Samples were incubated in 0.01% (w/v) aniline blue prepared in K□HPO□ buffer (pH 9.5) for 4 h at room temperature under gentle agitation in the dark, as described by Adam & Somerville (1996), with modifications. After staining, leaves were mounted on microscope slides in 50% (v/v) glycerol.

Callose deposits were visualized under UV excitation (365–405 nm) and quantified using an Olympus IX81 epifluorescence microscope (Olympus, Japan). Multiple images per leaf were acquired under identical acquisition settings across all samples for quantitative analysis. Callose deposition was quantified using Fiji by calculating the total fluorescent area after background subtraction, with the signal used as a proxy for callose accumulation. Representative images were acquired using a Leica TCS LSI confocal laser-scanning microscope (Leica Microsystems, Germany) under identical acquisition settings across samples and selected based on consistent staining patterns observed across biological replicates. For each biological replicate (n = 6; each replicate consisted of a single plant), two leaves were analyzed, and multiple images (4–7) were acquired per leaf using a standardized sampling approach that avoided trichomes and aphids. In total, 12 leaves per genotype and condition were analyzed.

### Gene expression analysis

For gene expression analysis, fully expanded leaves were harvested from 5-week-old *A. thaliana* plants (Col-0, *cslb4-1*, and *cslb4-2*) grown under basal (non-infested) conditions. Four biological replicates were analyzed (n = 4), each consisting of a single plant, from which two leaves were collected. Samples were immediately frozen in liquid nitrogen and subsequently ground to a fine powder using a TissueLyser II (Qiagen, Germany) with stainless steel beads in 2-mL tubes. Total RNA was extracted using TRIzol reagent, and genomic DNA contamination was removed by DNase I treatment. RNA integrity was verified by agarose gel electrophoresis, and concentration was determined spectrophotometrically. First-strand cDNA was synthesized from 1 µg of total RNA using the iScript cDNA Synthesis Kit (Bio-Rad, USA) according to the manufacturer’s instructions. Quantitative real-time PCR (RT–qPCR) was performed using Brilliant III Ultra-Fast SYBR Green Master Mix (Agilent Technologies, USA) in an AriaMx Real-Time PCR System (Agilent Technologies, USA). No-template controls and melt curve analyses were performed to verify amplification specificity. Primer sequences are listed in **Supplementary Table S2**.

Quantification cycle (Cq) values were obtained using AriaMx software. Amplification efficiencies (E) were estimated using LinRegPCR software (Untergasser *et al*., 2021), and the mean efficiency for each primer pair was used for relative quantification. Relative transcript abundance was calculated using an efficiency-corrected model (Pfaffl, 2001) and normalized using the geometric mean of two reference genes (Vandesompele *et al*., 2002). *TUB4* and *YLS8* were used as reference genes for normalization. Each reaction was performed in technical replicates to ensure reproducibility.

### Tissue enrichment and gene expression analysis

For tissue-specific gene expression analysis in vascular, mesophyll, and epidermal tissues, 30 adult aphids were placed on fully expanded leaves from 5-week-old *A. thaliana* Col-0 plants for 6 and 24 hours post-infestation (hpi), while leaves enclosed in empty cages were used as non-infested controls (0 h). Mesophyll, epidermal, and vascular tissues were isolated as previously described (Endo *et al*., 2016; Peppino Margutti *et al*., 2025) using a modified Tape–Arabidopsis Sandwich method, which enables the sequential separation of epidermal, mesophyll, and vascular tissues from the same leaf.

Tissue enrichment was validated by assessing the expression of tissue-specific marker genes, including *LHCB2.1* (mesophyll), *GC1* (epidermis), and *SULTR2.1* (vascular) (Endo *et al*., 2014) (**Supplementary Table S2; Supplementary Fig. S2**). Only samples showing clear enrichment of the corresponding markers were retained for further analysis.

For each biological replicate (n = 3), tissues were collected independently from pooled samples consisting of four leaves obtained from two independent plants (two leaves per plant). Isolated tissues were subsequently used for RNA extraction and RT–qPCR analysis as described above, and transcript levels were normalized to *YLS8* and expressed relative to whole-leaf expression levels for each gene and time point (0, 6, and 24 h).

### Subcellular localization of CSLB4

To determine the subcellular localization of CSLB4, the full-length coding sequence (CDS) of *A. thaliana CSLB4*, excluding the stop codon, was amplified from Col-0 cDNA and cloned in frame with GFP as a C-terminal fusion into the pGW405 binary vector using Gibson assembly under the control of the CaMV 35S promoter. Primer sequences used for cloning are listed in **Supplementary Table S2**. The resulting CSLB4–GFP construct was introduced into *Agrobacterium tumefaciens* strain GV3101 and transiently expressed in *Nicotiana benthamiana* leaf epidermal cells.

*Agrobacterium tumefaciens* cultures carrying CSLB4–GFP and subcellular markers for the Golgi (α-1,2-mannosidase-I–RFP) and endoplasmic reticulum (wall-associated kinase 2–RFP) (Nelson *et al.,* 2007) were grown overnight, harvested, and resuspended in infiltration buffer containing 10 mM MgCl□, 10 mM MES (pH 5.6), and 150 µM acetosyringone. Bacterial suspensions were adjusted to an OD[[□ of 0.8 and co-infiltrated into fully expanded leaves using a needleless syringe at a ratio of 1:3 (marker:CSLB4–GFP).

Fluorescence signals were examined 48 h post-infiltration using a confocal laser scanning microscope (Leica TCS SP8). GFP and RFP signals were acquired sequentially to avoid spectral overlap. GFP fluorescence was excited at 488 nm and detected at 500–550 nm, while RFP was excited at 561 nm and detected at 580–620 nm. Images were captured under identical acquisition settings for all samples. For each biological replicate (n = 3; each replicate consisted of an independently infiltrated leaf), multiple fields of view were imaged per sample. Experiments were repeated independently at least twice.

### Structural modeling and motif analysis

Structural models of CSLB3 (Q8RX83) and CSLB4 (O80891) were obtained from the SWISS-MODEL repository (Waterhouse *et al*., 2018). The CSLB4 model was retrieved as a predicted trimeric assembly consistent with the architecture of plant cellulose synthases. Structural superposition onto CesA7 (PDB ID: 7D5K) was performed to assess conservation of catalytic features (Zhang *et al*., 2021). To visualize predicted transmembrane regions, the CSLB4 model was embedded in a POPC lipid bilayer using the CHARMM-GUI server (Jo *et al*., 2008). Conserved motifs (DXD, DDG, TED, and QXXRW), associated with catalytic activity were identified based on previous annotations (Purushotham *et al*., 2020; Daras *et al*., 2021).

### Statistical analysis of experimental data

All statistical analyses were conducted in R (version 4.4.0) (R Core Team, 2024). Data distribution was assessed using the Shapiro–Wilk test and visual inspection of residuals. When necessary, phenotypic data were transformed to improve normality and homoscedasticity prior to analysis.

For normally distributed data, including no-choice assays, gene expression, and comparisons among haplotype classes, differences were evaluated using Welch’s t-test (two groups) or one-way analysis of variance (ANOVA). When comparing multiple groups against a control, post hoc comparisons were performed using Dunnett’s, and for all pairwise comparisons, Tukey’s HSD test was used. Callose quantification data were log-transformed prior to analysis and analyzed using a two-way ANOVA to assess the effects of genotype, aphid infestation, and their interaction. Gene expression in tissue-specific assays during aphid infestation was analyzed using the same two-way ANOVA framework, assessing the effects of genotype, tissue, and their interaction. When significant effects were detected, post hoc comparisons were performed using Tukey’s HSD test.

For EPG analyses, non-normally distributed variables were analyzed using the Wilcoxon rank-sum test (exact test where applicable). Proportional variables (%) were analyzed using chi-squared (χ^2^) tests. Differences were considered statistically significant at *p* < 0.05. All tests were two-sided. Specific statistical tests applied to each experiment are indicated in the corresponding figure legends.

## Results

### Genome-wide association mapping identifies loci associated with aphid resistance

To identify genomic regions underlying natural variation in resistance to the specialist aphid *Brevicoryne brassicae*, we performed a genome-wide association study (GWAS) using a panel of 200 *Arabidopsis thaliana* accessions from diverse ecological origins (Alonso-Blanco *et al*., 2016), spanning a broad geographic range (**Fig. 1A**). Principal component analysis (PCA) of the SNP revealed population structure consistent with the ten previously defined genetic groups in *A. thaliana* (Horton *et al*., 2012) (**Supplementary Fig. S3**). Aphid performance was assessed for each accession using a non-choice bioassay, in which resistance was measured as the total number of nymphs (offspring number) produced by two adult founder aphids (**Supplementary Fig. S1**). This analysis revealed phenotypic variation among accessions (**Fig. 1B**), while no significant differences in offspring number were observed among genetic groups (**Supplementary Fig. S3**), suggesting that population structure does not strongly influence phenotypic variation. The phenotype exhibited moderate broad-sense heritability (*H*^2^ = 0.41), indicating that a considerable proportion of phenotypic variation is genetically determined. Together, these results establish that resistance to *B. brassicae* is a quantitative trait shaped by natural genetic variation, supporting the use of GWAS to identify underlying loci.

**Figure 1.**
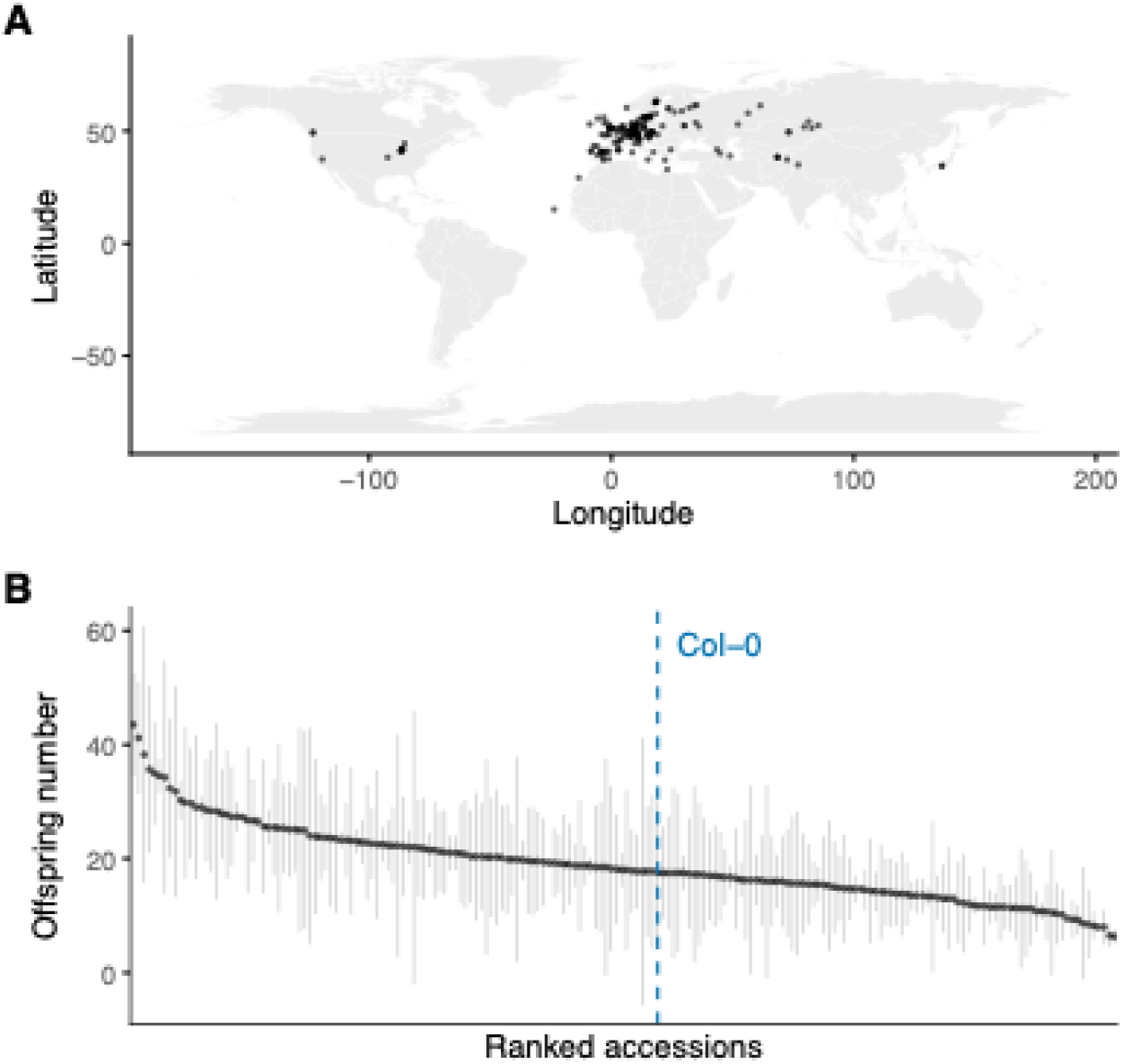
Natural variation in aphid performance of *B. brassicae* across *A. thaliana* accessions. **(A)** Geographic distribution of the 200 *A. thaliana* accessions included in the genome-wide association study (GWAS), capturing broad geographic and ecological diversity. **(B)** Phenotypic variation in aphid performance across accessions, measured as the total offspring number per accession. Plants were infested with two immature female nymphs (founders), and the offspring number was recorded at 19 days post-infestation. Points represent accession means, and error bars indicate standard error. The dotted line indicates the position of the Col-0 accession.

GWAS identified a single SNP surpassing the Bonferroni-corrected significance threshold (SNP1), along with two additional loci on different chromosomes showing suggestive association signals (SNP2 and SNP3; -log_10_(*p*) > 7; **Fig. 2A; Supplementary Table S3**). These three regions were further investigated through haplotype-based analysis to assess the combined effects of linked variants.

**Figure 2.**
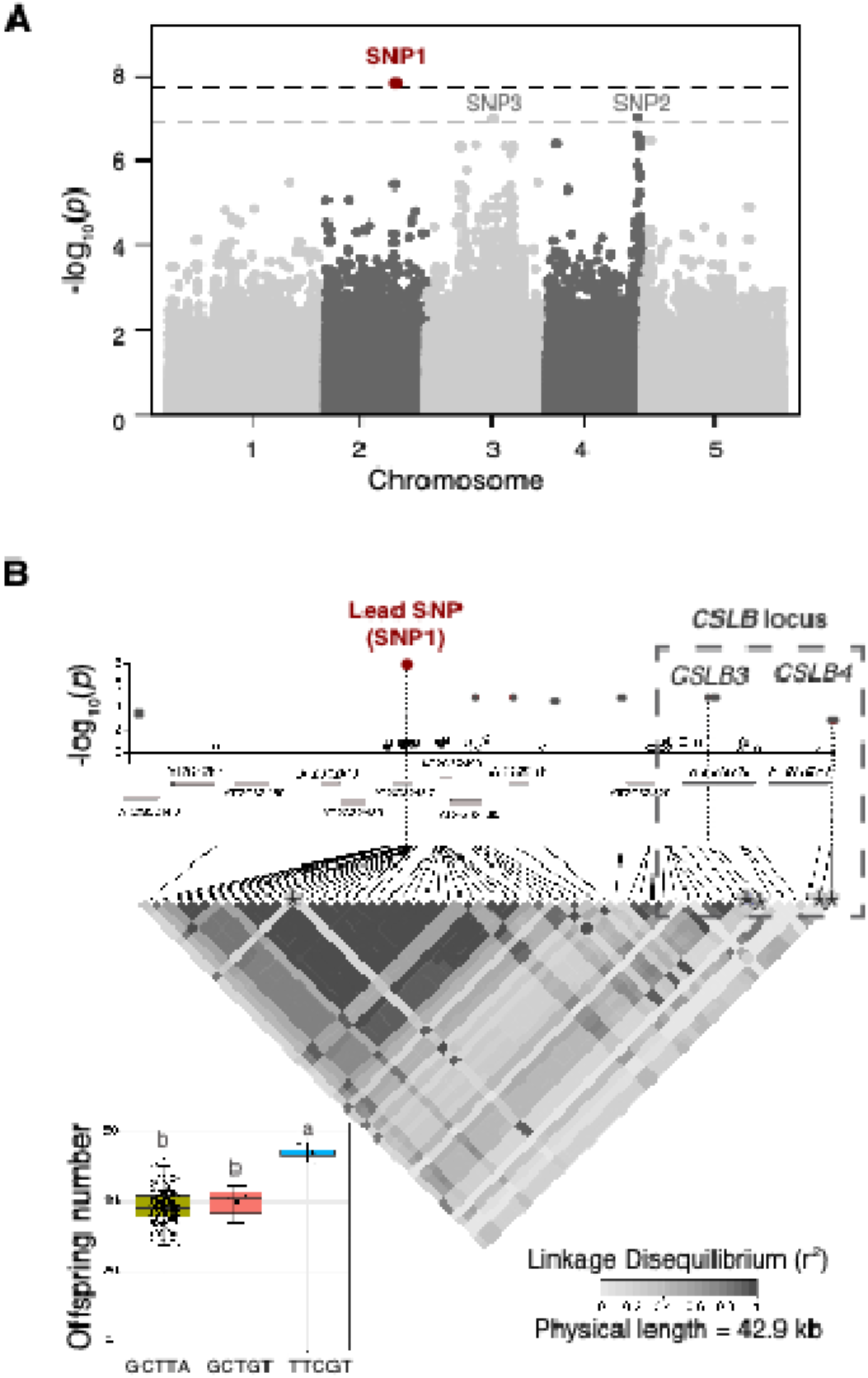
Genome-wide association mapping identifies a *CSLB*-containing locus underlying natural variation in aphid performance. **(A)** Manhattan plot showing the association between SNP markers and aphid performance of *B. brassicae* across the five *A. thaliana* chromosomes. Association testing was performed using a mixed linear model that accounts for population structure. The lead SNP (SNP1), surpassing the Bonferroni-corrected threshold, is highlighted, while SNP2 and SNP3 represent suggestive association signals. Each point represents a single SNP. The upper dashed line indicates the Bonferroni-corrected genome-wide significance threshold, while the lower dashed line represents a suggestive significance threshold (-log_10_(*p*) > 7). **(B)** Local association plot of the most significant locus identified on chromosome 2. The lead SNP (SNP1) is highlighted in red, and surrounding SNPs are colored according to their level of linkage disequilibrium (LD; r²). Gene models within the associated interval are shown below, including two members of the *Cellulose Synthase-Like B* (*CSLB*) family, *CSLB3* and *CSLB4*. The LD heatmap depicts the local haplotype structure across the region. Asterisks indicate SNPs included in the haplotype analysis. The inset boxplot shows aphid performance across allelic classes at the lead SNP, revealing significant differences in offspring numbers among haplotypes. These results identify a *CSLB*-containing locus that contributes to natural variation in aphid performance.

Haplotype analysis revealed significant differences in aphid performance for SNP1 and SNP2 (**Table 1**), whereas no consistent haplotypes associated were detected for the SNP3. The locus harboring the strongest association signal (SNP1) showed the most robust and biologically consistent effects and was therefore selected for detailed characterization. To fine-map this locus, linkage disequilibrium (LD) was examined within the SNP1 region. The lead SNP was in significant LD (r^2^ > 0.1, *p* < 0.001) with 12 neighboring SNPs, of which eight also showed elevated association signals (-log_10_ > 4; **Fig. 2B**). Four of these SNPs mapped within the *CSLB3* (AT2G32530) and *CSLB4* (AT2G32540) gene regions. Based on their genomic proximity, strong signals, and LD with the lead SNP (SNP1), haplotypes were constructed by combining these SNPs with the lead variant. Three informative haplotypes showed significant differences in aphid performance, with accessions carrying the TTCGT haplotype exhibiting significantly higher offspring number (**Fig. 2B**), consistent with increased susceptibility. These results indicated that the association signal likely reflects the combined effects of linked variants within this locus rather than a single causal polymorphism. The presence of associated SNPs in both *CSLB* genes, together with their close genomic proximity, suggests a potential shared or partially redundant contribution to the observed phenotypic variation. However, the extent to which each gene contributes to aphid performance remains difficult to disentangle based on LD and haplotype structure alone.

**Table 1.** Candidate genes identified by haplotype analysis within genomic regions defined by GWAS-associated lead SNPs. Each gene is assigned to its corresponding lead SNP (SNP1–SNP3; see Fig. 2A). Haplotypes are denoted as H1–Hn.

| Gene | Chr | GWAS SNP | Description | GSS* | p-value | q-value | Haplotypes |
| --- | --- | --- | --- | --- | --- | --- | --- |
| AT2G32440 | 2 | SNP1 | Ent-kaurenoic acid hydroxylase | 12.79 | 0.00167 | 0.02895 | H1–H3 |
| AT2G32487 | 2 | SNP1 | Putative protein (unknown function) | 30.35 | 0.00001 | 0.00059 | H1–H5 |
| AT2G32500 | 2 | SNP1 | Stress-responsive alpha/beta barrel domain protein | 22.22 | 0.00111 | 0.02199 | H1–H6 |
| AT2G32510 | 2 | SNP1 | MEKK subfamily protein involved in wound- and JA-induced signaling | 19.18 | 0.00178 | 0.02962 | H1–H5 |
| AT2G32520 | 2 | SNP1 | Alpha/beta-Hydrolases superfamily protein | 22.56 | 0.00015 | 0.00347 | H1–H5 |
| AT2G32530 | 2 | SNP1 | Cellulose synthase-like protein | 21.54 | 0.00025 | 0.00520 | H1–H4 |
| AT2G32540 | 2 | SNP1 | Cellulose synthase-like protein | 23.57 | 0.00003 | 0.00104 | H1–H4 |
| AT4G37440 | 4 | SNP2 | Putative protein (unknown function) | 26.72 | 0.00002 | 0.00083 | H1–H5 |
| AT4G37445 | 4 | SNP2 | Calcium ion-binding protein | 27.94 | 0.00004 | 0.00128 | H1–H6 |
| AT4G37450 | 4 | SNP2 | Lysine-rich arabinogalactan-protein (AGP) | 24.04 | 0.00001 | 0.00059 | H1–H3 |
| AT4G37460 | 4 | SNP2 | Tetratricopeptide repeat (TPR) domain protein | 25.12 | 0.00001 | 0.00059 | H1–H3 |
| – | 3 | SNP3 | No significant haplotypes associations detected | – | – | – | – |

\* GSS, global score statistic. Chr, chromosome
Corresponding haplotype classes and sequences are provided in Supplementary Table S4.
Lead SNP positions (TAIR10): SNP1 (Chr2: 13,792,857 bp), SNP2 (Chr4: 17,603,749 bp), SNP3 (Chr3: 13,002,652 bp).

Given that both *CSLB3* and *CSLB4* are located within the associated haplotype block (**Fig. 2B**), both genes were considered as potential contributors to aphid performance variation. However, publicly available RNA-seq data visualized using the eFP Browser (ePlant; **Supplementary Fig. S4**) (Klepikova *et al*., 2016) indicate that *CSLB3* displays a relatively broad expression pattern across tissues, whereas *CSLB4* shows higher expression, particularly in leaves. To further assess the relevance of both *CSLB* genes during aphid infestation, we analyzed their temporal expression patterns and tissue-specific expression in leaves. To ensure the specificity of the isolated tissue fractions, enrichment was validated using tissue-specific marker genes (**Supplementary Fig. S2**). *CSLB4* transcript levels increased specifically in epidermal tissues at 24 h post-infestation, whereas *CSLB3* did not show significant changes (**Fig. 3A**). Because the epidermis represents the initial site of aphid stylet penetration and feeding, this tissue-specific induction provides additional support for prioritizing *CSLB4* for functional analysis in the context of aphid interactions. To gain insight into the potential biochemical function of CSLB proteins, we performed sequence and structural analyses of CSLB3 and CSLB4. Structural modeling suggested that CSLB proteins adopt a fold similar to that of the cellulose synthase complex (**Supplementary Fig. S5**). Consistent with their classification within the glycosyltransferase family 2 (GT2), this supports a role in the biosynthesis of non-cellulosic cell wall (CW) polysaccharides. Also, CSLB3 and CSLB4 share high sequence identity (85.4%), with conservation concentrated in the predicted GT2 catalytic domain and associated functional regions (**Supplementary Fig. S5**). Notably, conserved regions include catalytic motifs characteristic of, as well as regions predicted to form the substrate-binding pocket (**Fig. 3B**). Despite their high sequence similarity, *CSLB3* and *CSLB4* display distinct tissue-specific expression patterns and exhibit subtle differences within the predicted catalytic domain, suggesting potential functional divergence. Accordingly, *CSLB4* was selected for functional validation based on its position within the associated locus, its tissue-specific induction during aphid infestation, and its predicted involvement in CW-related processes. Together, these results identified *CSLB4* as a strong candidate gene underlying natural variation in aphid resistance.

**Figure 3.**
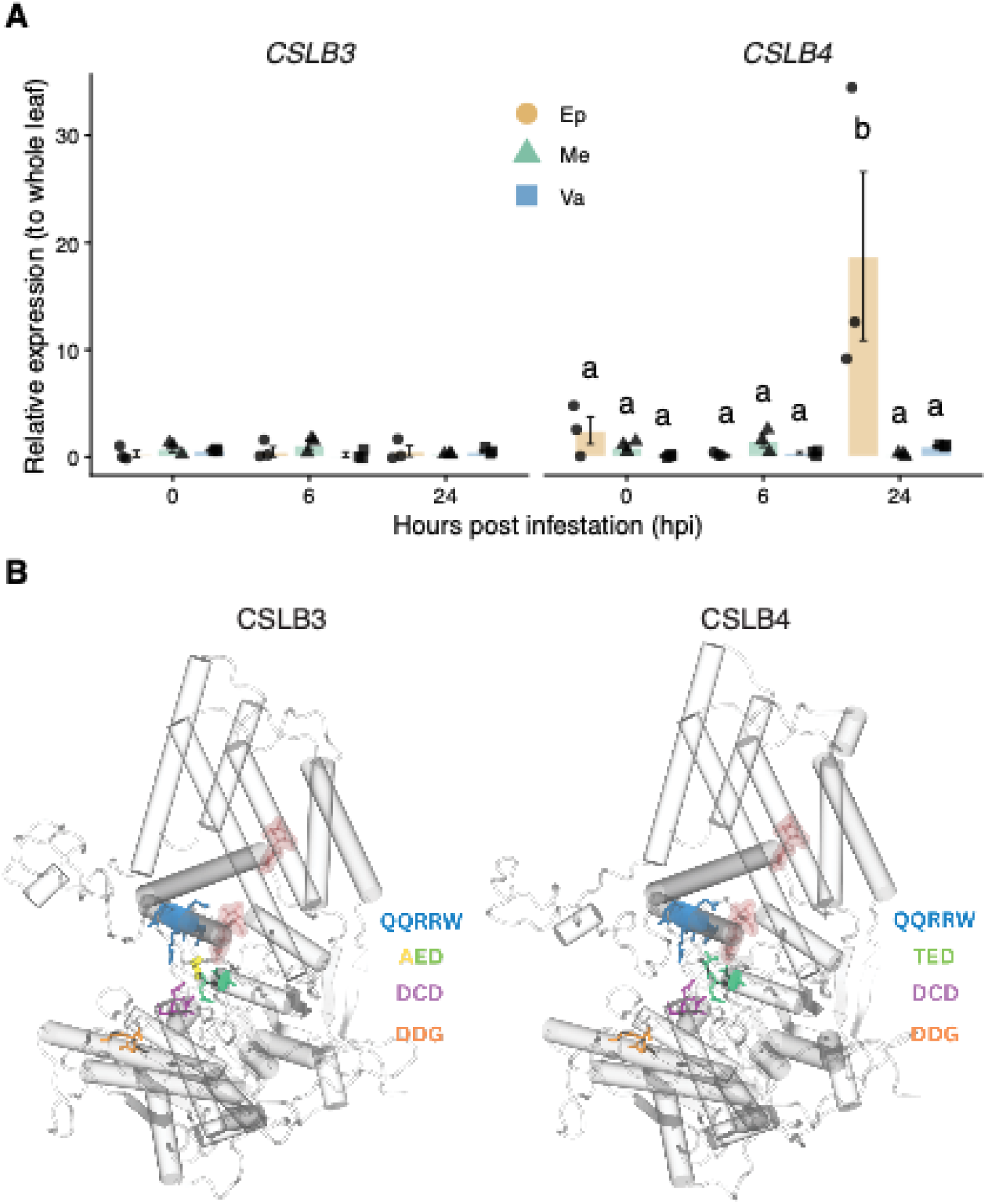
Spatio-temporal expression and predicted structural features of CSLB3 and CSLB4. **(A)** Transcript levels in epidermis (Ep), mesophyll (Me), and vascular tissues (Va) at 0, 6, and 24 hours post-infestation (hpi). Expression values are relative to whole-leaf levels and normalized to *YLS8*, reflecting the relative contribution of each tissue to total transcript accumulation. Bars represent mean ± standard error; points represent biological replicates (n = 3 per condition; each replicate consisted of four leaves from two independent 5-week-old plants). Different letters indicate significant differences among tissues (Ep, Me, Va) within each time point (Tukey’s HSD test, *p* < 0.05). **(B)** Structural monomeric models of CSLB3 and CSLB4, highlighting conserved catalytic motifs and the predicted substrate-binding pocket. The native β-cellobiose ligand from CesA7 is shown in red as a positional reference for the predicted binding site.

### Disruption of CSLB4 alters aphid performance

To assess the functional relevance of *CSLB4*, we analyzed aphid performance using loss-of-function mutant plants. Two independent T-DNA insertion lines, *cslb4-1* and *cslb4-2*, with insertions in distinct regions of the *CSLB4* gene (**Fig. 4A**), showed a strong reduction in *CSLB4* transcript compared with WT, confirming effective gene disruption (**Fig. 4B**). We then evaluated aphid performance using a no-choice assay with *B. brassicae*. Both *cslb4* mutant lines supported significantly fewer aphid offspring compared with WT (**Fig. 4C**), showing reductions of 56.3% (*cslb4-1*) and 39.2% (*cslb4-2*). The consistent phenotype observed across two independent alleles supports that this effect is associated with loss of *CSLB4* function. Together, these results demonstrate that *CSLB4* contributes to plant susceptibility to the *B. brassicae*, as its disruption leads to reduced aphid reproductive success.

**Figure 4.**
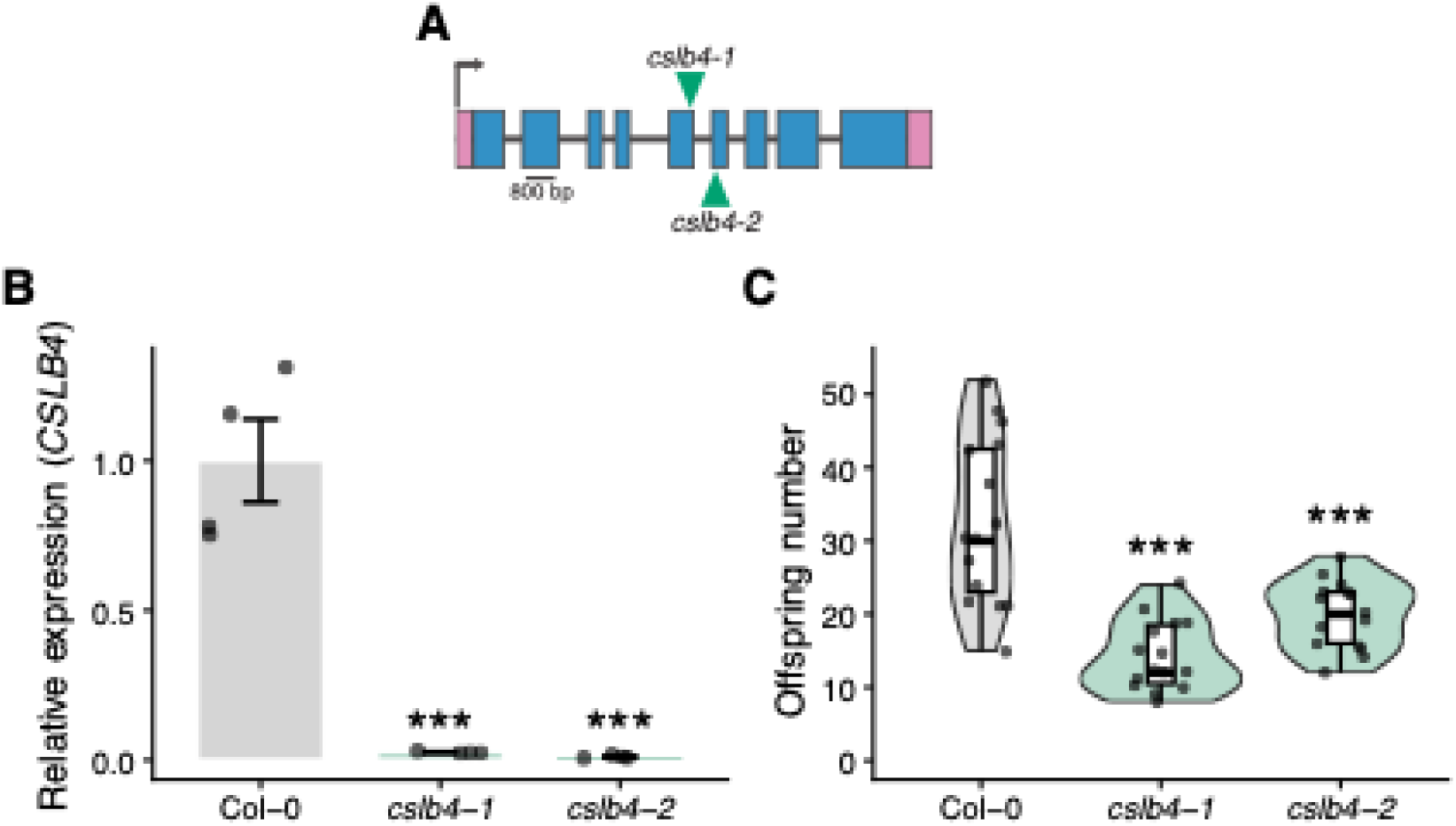
Loss of *CSLB4* reduces aphid performance. **(A)** Gene structure of *CSLB4* showing the positions of T-DNA insertions in the *cslb4-1* and *cslb4-2* lines. Boxes represent exons, pink boxes indicate untranslated regions (UTRs), and lines represent introns. Green triangles mark the positions of T-DNA insertions. Scale bar = 800 bp. **(B)** Relative transcript levels of *CSLB4* in Col-0 and the two mutant lines. Expression was normalized to reference genes and expressed relative to Col-0. Bars represent mean ± standard error (n = 4; each replicate consisted of a single plant, from which two leaves were collected). **(C)** Non-choice assay of aphid performance, measured as the total number of offspring produced per plant at 19 days post-infestation of 5-week-old Col-0, *cslb4-1*, and *cslb4-2* with *B. brassicae*. Violin plots represent the distribution of biological replicates (n = 15; each replicate consists of a single plant infested with two founder aphids). Boxes indicate the interquartile range (IQR), horizontal lines represent the median, whiskers extend to 1.5 × IQR. Dots correspond to individual data points. Statistical differences in (B) and (C) were assessed using a one-way ANOVA followed by Dunnett’s multiple comparisons test against Col-0. Asterisks indicate significant difference relative to Col-0 genotypes (***, *p* < 0.001).

Given the predicted role of *CSLB4* in CW–related processes, and evidence that the plant CW functions not only as a structural barrier but also as a dynamic regulator of plant–microbe and plant–aphid interactions (Silva-Sanzana *et al*., 2020; Molina *et al*., 2024b), we next asked whether its effect on aphid performance was specific to the specialist aphid *B. brassicae* or extended to other aphid species. To address this, we assessed aphid performance on *cslb4-1* using the generalist aphid *Myzus persicae*. No significant differences in offspring number were detected between *cslb4-1* and WT (**Supplementary Fig. S6**), indicating that disruption of *CSLB4* does not confer broad resistance to aphids. Together, these results show that *CSLB4* affects aphid performance in a species-specific manner, reducing offspring production in *B. brassicae* but not in *M. persicae*, and suggest a role in modulating host traits associated with CW properties.

### CSLB4 localizes to the Golgi apparatus

To gain insight into the cellular functions of *CSLB4*, we examined its subcellular localization. The coding sequence of *A. thaliana CSLB4* was fused to GFP and transiently expressed in *Nicotiana benthamiana* leaf epidermal cells. CSLB4–GFP displayed discrete punctate cytoplasmic structures (**Fig. 5**). Co-expression with a Golgi–RFP marker revealed substantial co-localization of fluorescent signals in merged images, indicating that CSLB4 localizes to Golgi-associated compartments. In contrast, no co-localization was observed with an endoplasmic reticulum (ER) marker, supporting the specificity of Golgi localization.

**Figure 5.**
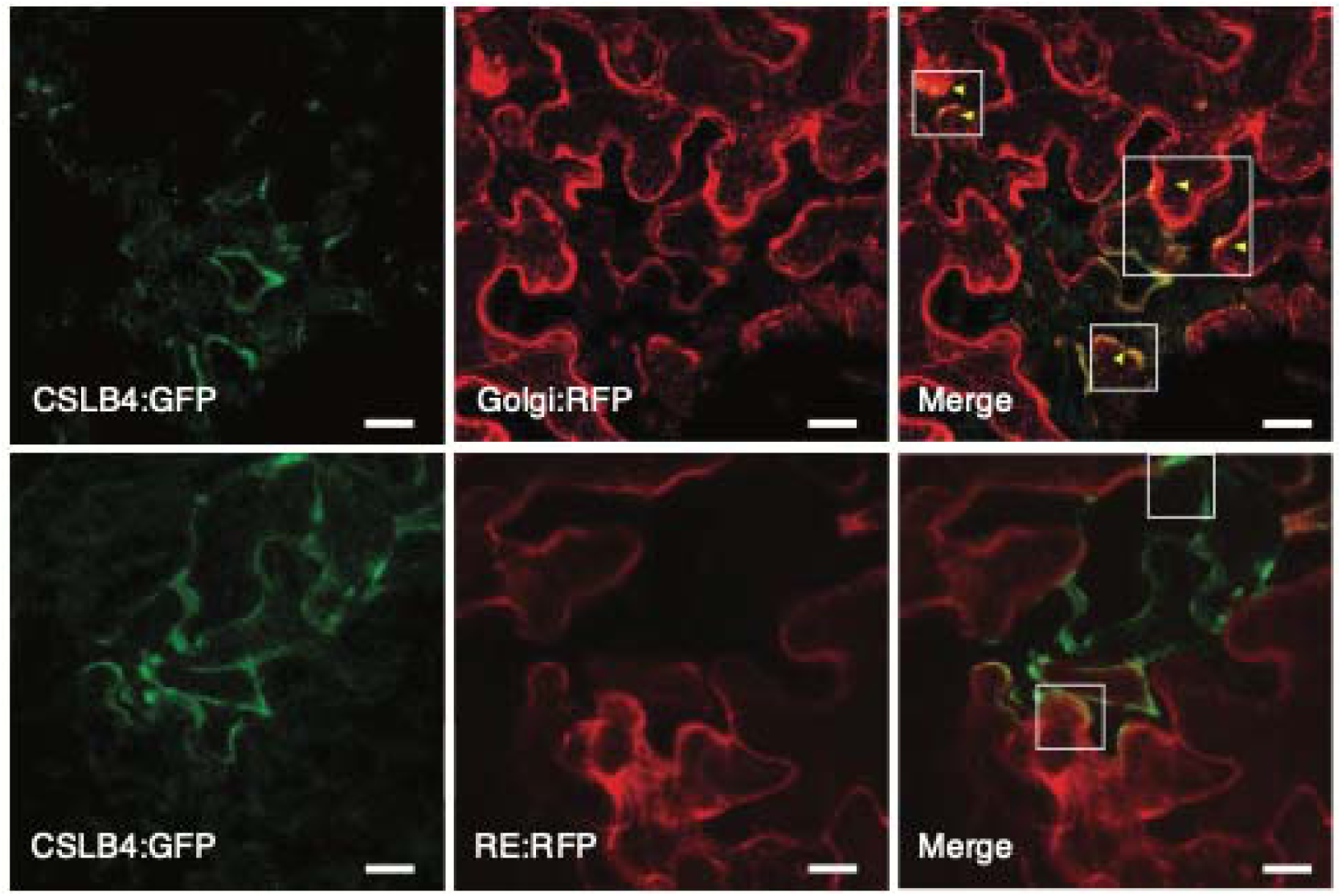
Subcellular localization of CSLB4 in *N. benthamiana* leaf epidermal cells. Confocal images of CSLB4–GFP transiently expressed in *N. benthamiana* leaf epidermal cells. CSLB4–GFP signal is observed in discrete cytoplasmic puncta. Co-expression with a Golgi marker (α-1,2-mannosidase-I–RFP) and an endoplasmic reticulum marker (wall-associated kinase 2–RFP) shows partial overlap of fluorescence signals in merged images. Insets highlight regions of signal overlap. Images are representative of independent biological replicates. Scale bars, 20 μm.

This localization is consistent with the reported distribution of cellulose synthase-like (CSL) proteins, members of the GT2 family (Cantarel *et al*., 2009), which are typically associated with the Golgi apparatus and involved in the biosynthesis of non-cellulosic polysaccharides such as hemicelluloses (Liepman *et al*., 2005; De Caroli *et al*., 2014). Together with the predicted structural features of CSLB4 (**Fig. 3B**), these findings support a role for CSLB4 in CW polysaccharide biosynthesis or remodeling.

### Cell wall composition and organization are altered in cslb4 mutants

Given the predicted role of CSLB4 in CW–related processes and its Golgi localization, we therefore examined whether loss of *CSLB4* affects CW composition. To obtain an integrated view of CW-associated biomass and polysaccharide composition, we analyzed alcohol-insoluble residues (AIR), which represents the bulk CW fraction. AIR analyses revealed a significant decrease in galactose, galacturonic acid (GalA), and arabinose content in *cslb4-1* compared with WT, with GalA showing a marked reduction (54.1%; **Fig. 6A**). No significant differences were detected for the remaining monosaccharides. Given that GalA is a major component of pectin, these results suggest that pectic polysaccharides are altered in *cslb4-1*. Together with the decreased levels of Gal and Ara, these results are consistent with modifications in matrix polysaccharides, potentially reflecting changes in the composition or side-chain decoration of non-cellulosic CW polysaccharides.

**Figure 6.**
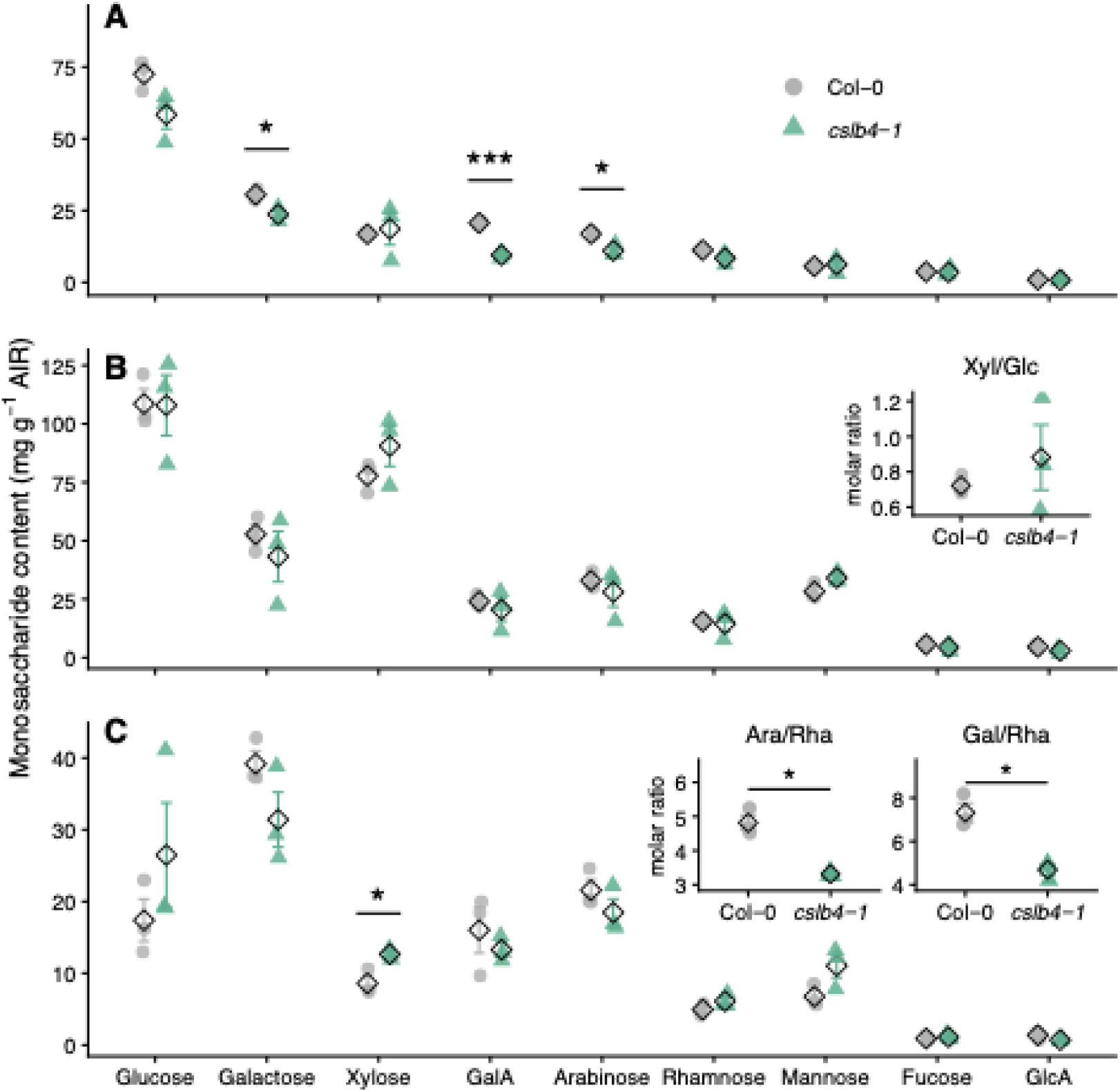
Loss of *CSLB4* alters cell wall monosaccharide composition. Alcohol-insoluble residue (AIR) was extracted from 5-week-old Col-0 and *cslb4-1*. Monosaccharide content (mg g^-1^ AIR) was determined by HPAEC–PAD in **(A)** total AIR, **(B)** hemicellulose-enriched fraction, including the Xyl/Glc molar ratio, and **(C)** pectin-enriched fractions, including diagnostic rhamnogalacturonan I (RG-I) molar ratios (Ara/Rha and Gal/Rha). Points represent biological replicates (n = 3; each replicate consisted of pooled rosettes from six plants), and diamonds indicate the mean ± standard error. Asterisks indicate significant differences between genotypes (Student’s t-test: \**p* < 0.05, \*\*\**p* < 0.001).

To further resolve these changes, AIR was fractionated into hemicellulose- and pectin-enriched fractions. No differences in monosaccharide composition or Xyl/Glc molar ratio were observed in the hemicellulose-enriched fractions (**Fig. 6B**). To assess potential alterations in xyloglucan (XyG) structure not captured by monosaccharide composition, immunodot blot analyses were performed using three XyG-specific antibodies: LM15 (XXXG), LM24 (XLLG), and LM25 (XXXG, XLLG, XXLG) (**Supplementary Fig. S7A,B**). No differences in antibody binding were detected, indicating that the *cslb4-1* mutation does not affect hemicellulose composition or XyG structure.

In contrast, analysis of pectin-enriched fractions revealed a significant increase in xylose content, likely reflecting minor hemicellulose carryover during fractionation (**Fig. 6C**). Although no differences were observed in the major pectic monosaccharides, significant changes were detected in specific monosaccharide ratios (**Fig. 6C**), including reduced arabinose-to-rhamnose (Ara/Rha) and galactose-to-rhamnose (Gal/Rha) ratios in *cslb4-1*. These ratios are commonly used as indicators of rhamnogalacturonan I (RG-I) side-chain composition, suggesting targeted changes in pectic architecture. In addition, methanol content was markedly reduced in *cslb4-1*, consistent with a lower degree of pectin methylesterification, compared to WT (**Supplementary Fig. S7C**). Together, these results indicate that loss of *CSLB4* is associated with specific alterations in pectin composition and architecture, with minimal effects on hemicellulose.

Given the absence of major changes in hemicellulose monosaccharide composition, we next assessed whether *CSLB4* affects hemicellulose features within the CW. Because XyG is the predominant hemicellulosic polysaccharide in the primary cell walls of Arabidopsis leaves (Zablackis *et al*., 1995), we used XyG epitope distribution and accessibility as a proxy for potential changes in hemicellulose organization. XyG epitope distribution and accessibility were assessed by immunofluorescence using the LM25 antibody. Leaf cross-sections revealed increased LM25 signal intensity in *cslb4-1* compared with WT (**Fig. 7A; Supplementary Fig. S8**). Quantitative image analysis using Fiji confirmed higher LM25 signal in mesophyll cell walls of *cslb4-1*, whereas no differences were observed in epidermal cell walls (**Fig. 7B**). Importantly, this increase occurs despite unchanged overall XyG epitope abundance between genotypes. These findings indicate that loss of *CSLB4* alters XyG epitope accessibility and/or spatial distribution within the CW, with increased LM25 signal specifically in mesophyll cell walls despite unchanged hemicellulose monosaccharide composition, consistent with changes in polysaccharide organization rather than total epitope abundance. Together, these results support a role for CSLB4 in modulating CW architecture by affecting polysaccharide organization rather than overall CW composition.

**Figure 7.**
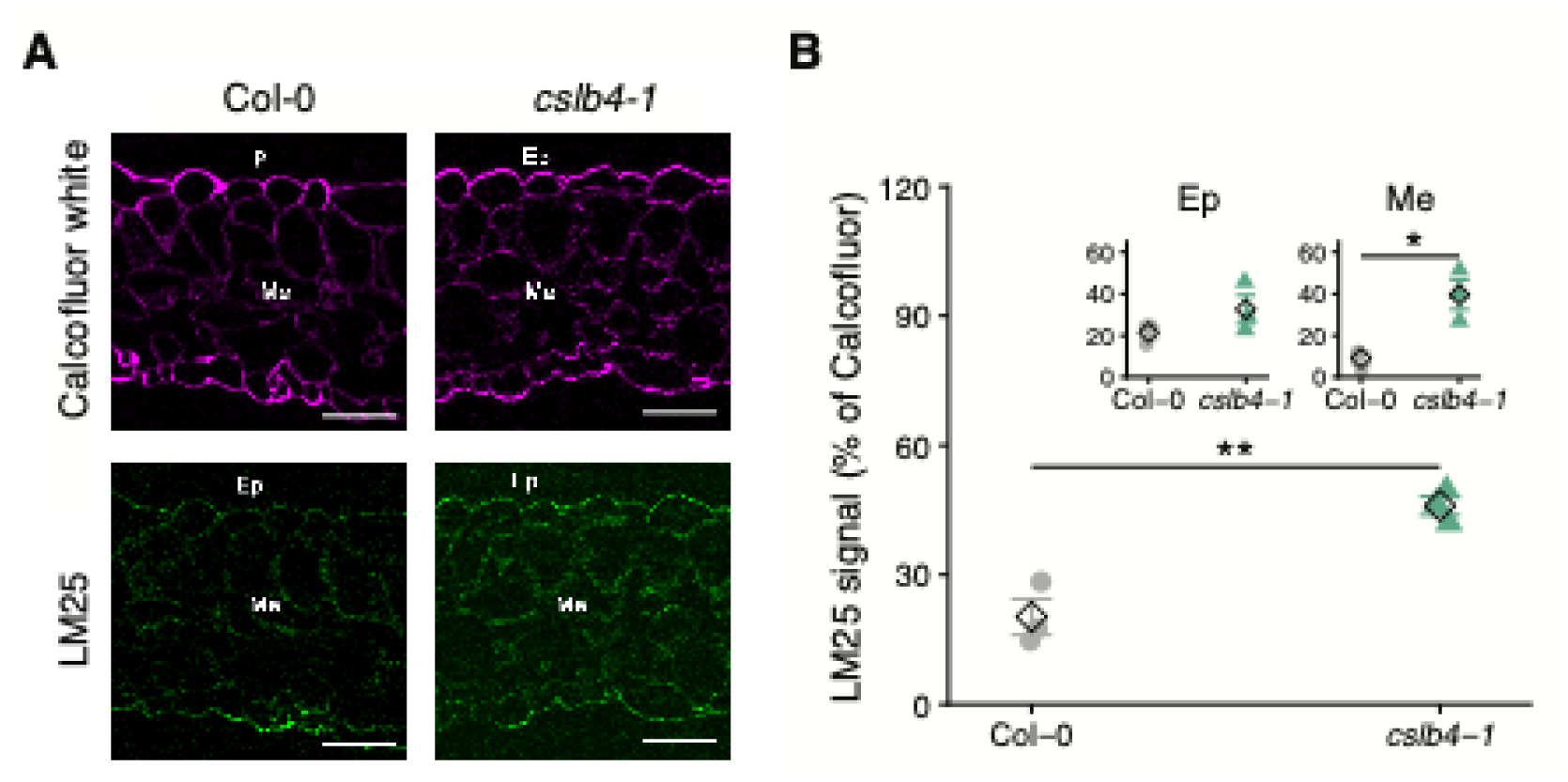
Loss of *CSLB4* alters xyloglucan distribution in leaf tissues. **(A)** Confocal images of leaf cross-sections from 5-week-old Col-0 and *cslb4-1* immunolabeled with the LM25 antibody, which recognizes xyloglucan (XyG) epitopes (XXXG, XXLG, and XLLG) (green). Calcofluor white was used to label cell walls (magenta). Images show representative cross-sections (n = 3 biological replicates), highlighting mesophyll (Me) and epidermal (Ep) cells. Additional images are provided in Supplementary Fig. S7. Scale bars, 50 μm. **(B)** Quantification of LM25 fluorescence intensity in mesophyll and epidermal cell walls using Fiji. Fluorescence intensity was normalized to the Calcofluor white signal to account for differences in cell wall content. Points represent biological replicates (n = 3; each replicate consisted of a single plant, from which two leaves were collected, for sectioning and imaging), and diamonds indicate the mean ± standard error. Asterisks indicate significant differences between genotypes (Student’s t-test: \**p* < 0.05, \*\**p* < 0.01).

### Aphid-feeding behavior is altered in cslb4-1

Given the observed changes in CW composition and architecture in *cslb4-1*, we next sought to determine whether these alterations affect aphid-feeding behavior. Because aphid stylets navigate through the apoplast and directly interact with the CW matrix, changes in polysaccharide organization could influence stylet progression and feeding dynamics (Silva-Sanzana *et al*., 2020). To test this, we performed electrical penetration graph (EPG) recordings, which measure changes in electrical resistance as the aphid stylet penetrates plant tissues, enabling discrimination of distinct probing and feeding phases, to characterize aphid probing and phloem feeding behavior on WT and *cslb4-1*.

EPG analysis revealed that feeding behavior of *B. brassicae* is significantly altered in *cslb4-1* across multiple phases of stylet activity (**Table 2**). During the pre-phloem phase, aphids on *cslb4-1* showed a longer duration of the first probe and a reduced proportion of time spent in pathway activities (C phase), along with an increased frequency of intracellular punctures (pd), indicating enhanced intracellular sampling during tissue exploration. In addition, aphids required fewer probes before reaching the first phloem ingestion event (E2). Consistent with these observations, aphids reached phloem-related phases significantly faster in *cslb4-1*, as indicated by a reduced time to first E and first E2 (**Table 2**). Despite this accelerated access to phloem, aphids on *cslb4-1* exhibited marked changes in phloem feeding behavior, including an increased number and duration of salivation events (E1) and a higher E2/C ratio, indicative of altered feeding dynamics after vascular contact.

**Table 2.**
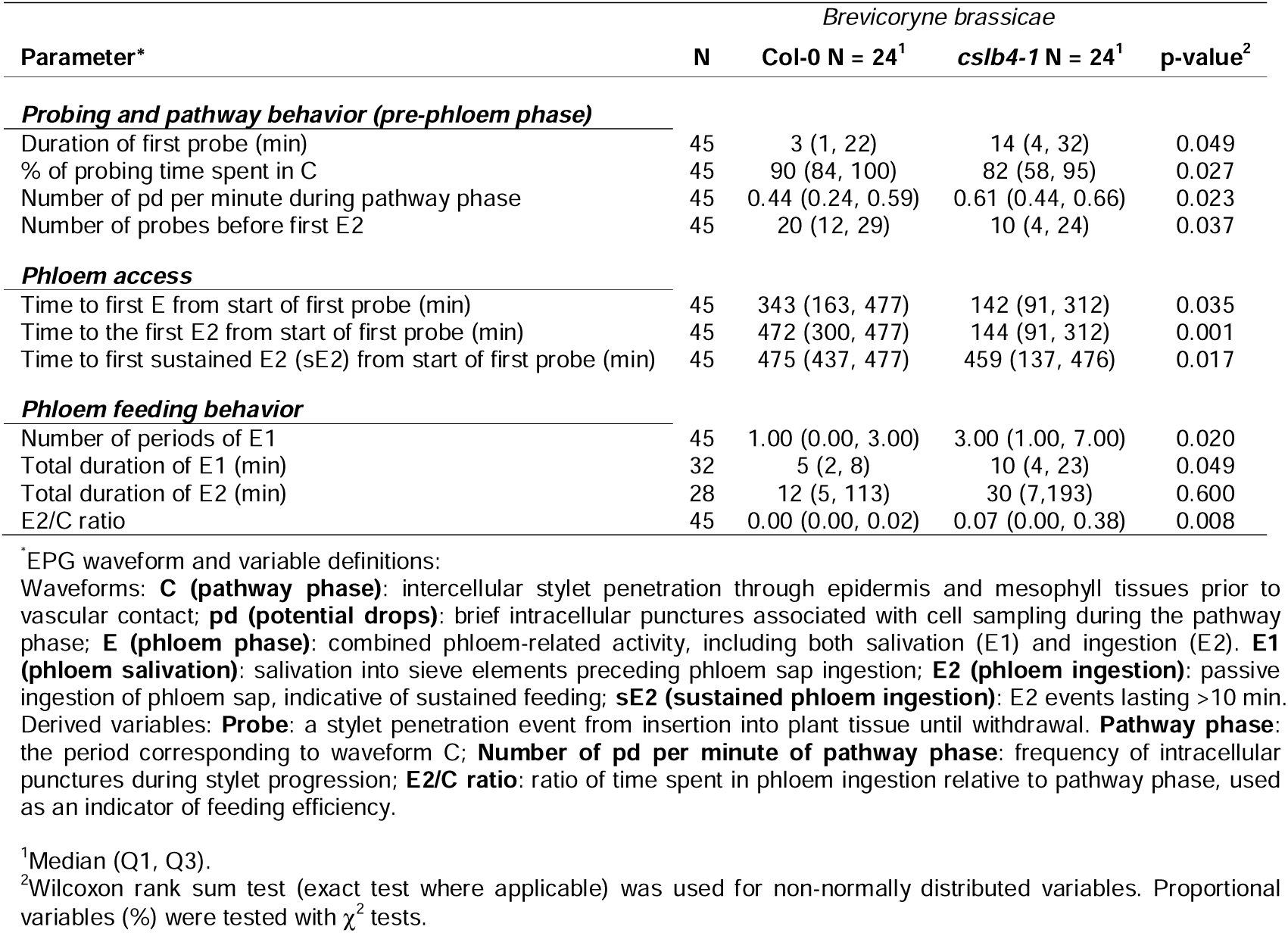
Electrical penetration graph (EPG) parameters describing feeding behavior of *B. brassicae* on Col-0 and *cslb4-1* plants during 8-h recordings.

Collectively, these parameters indicate that loss of *CSLB4* function does not impair aphid access to the phloem but instead alters both early probing behavior and feeding dynamics after vascular contact (**Table 2**). The combination of faster phloem access with altered salivation and ingestion behavior suggests that aphid performance is not limited by resource accessibility per se but rather by processes that affect feeding efficiency or host suitability.

These results reveal a decoupling between phloem accessibility and aphid performance, highlighting the importance of pre- and post-phloem processes in determining resistance. Together, these findings support a model in which CSLB4-dependent changes in CW architecture modify the structural and biochemical environment encountered by the stylet, thereby altering feeding dynamics without restricting phloem accessibility. This is further consistent with the epidermis-specific induction of CSLB4, suggesting a potential role in modulating cell wall properties at early stages of stylet penetration.

### Loss of CSLB4 affects callose deposition in response to aphid feeding

Given that CSLB4-dependent changes in CW composition and architecture alter aphid feeding behavior, as revealed by EPG analyses (**Table 2**), we next examined whether these modifications also impact callose deposition, a major CW reinforcement response at aphid feeding sites (Will and van Bel, 2006). Callose deposition is a well-characterized basal response during biotic interactions (Voigt, 2014) and is often associated with local CW perturbations caused by aphid stylet activity, although it also participates in multiple physiological processes (Li *et al*., 2023). To assess this, we quantified callose deposition in leaves following aphid infestation. In non-infested plants, callose levels were low and did not differ between WT and *cslb4-1* (**Fig. 8A; Supplementary Fig. S9**). However, after infestation with *B. brassicae*, WT plants showed increased callose deposition, whereas *cslb4-1* displayed a significantly reduced callose response compared to WT. Quantitative image analysis confirmed a reduced callose-positive area (measured as fluorescence pixel area) in *cslb4-1* leaves following aphid infestation (**Fig. 8B**). Consistent with these observations, no significant genotype × infestation interaction was detected, whereas a significant main effect of genotype indicated consistently lower callose deposition in *cslb4-1* compared with WT across treatments.

**Figure 8.**
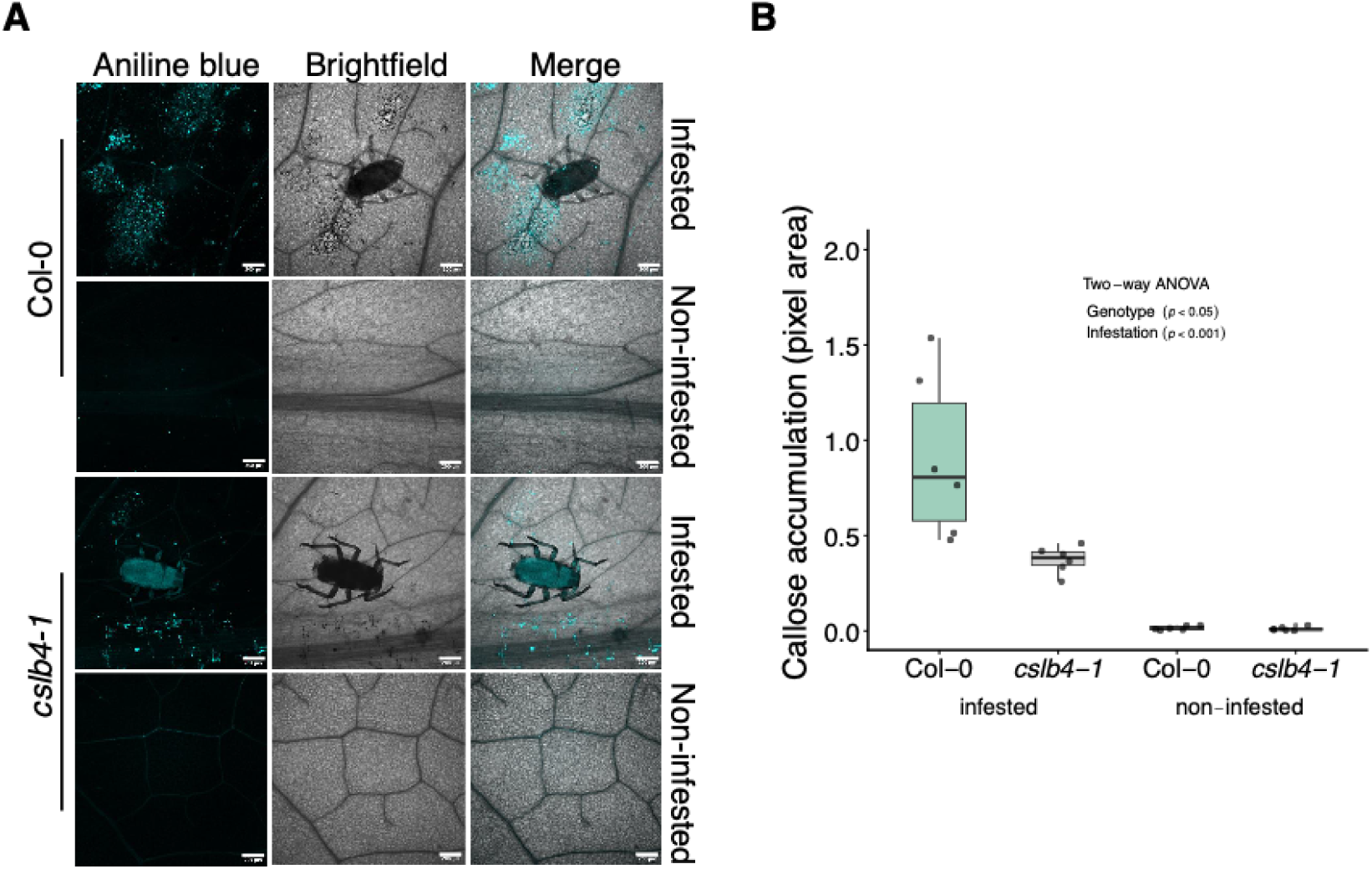
Loss of *CSLB4* reduces callose deposition in response to aphid infestation. Leaves from 5-week-old Col-0 and *cslb4-1* plants, either non-infested or infested with *B. brassicae* for 48 h, were stained with aniline blue to visualize callose deposition. **(A)** Confocal images of leaf cross-sections showing callose deposits (cyan). Merged images of aniline blue fluorescence and brightfield are shown. Scale bars, 200 μm. Images are representative of six biological replicates (additional images are provided in Supplementary Fig. S8). **(B)** Callose area was quantified from 4–7 images per leaf across six biological replicates per treatment (n = 6; 12 leaves per genotype and condition). Box plots show the interquartile range (IQR) with the median indicated by a horizontal line; whiskers extend to 1.5 × IQR. Two-way ANOVA detected significant effects of genotype (*p* < 0.05) and infestation (*p* < 0.001), with no significant genotype × infestation interaction.

These results indicate that loss of *CSLB4* reduces callose deposition, consistent with altered CW–associated responses. Despite this reduction, aphid reproductive performance was also decreased in the mutant background (**Fig. 4**), indicating that callose deposition is unlikely to be the primary determinant of resistance in this system. Instead, these findings suggest that CSLB4 influences CW-associated processes that contribute to the plant response to aphid feeding, with callose deposition representing only one component. Together with the observed changes in CW composition and architecture, these results support a model in which enhanced resistance in *cslb4*-1 is associated with broader alterations in CW architecture rather than a single defense response.

## Discussion

In this study, we identified CSLB4 as an uncharacterized modulator of aphid resistance through an integrative approach combining natural variation, functional genetics, and physiological analyses. Using GWAS, we uncovered a locus associated with aphid performance that includes *CSLB* genes and further demonstrated that loss of *CSLB4* function reduces aphid performance and host preference in Arabidopsis. Mechanistically, our results suggest that *CSLB4* disruption leads to specific alterations in cell wall (CW) composition and architecture, particularly affecting pectic components and xyloglucan organization, thereby modifying aphid-feeding behavior without restricting phloem access. These findings reveal a decoupling between phloem accessibility and aphid performance, highlighting the importance of CW architecture in determining plant–aphid interactions.

### Natural variation identifies cell wall–associated determinants of aphid resistance

GWAS provide a powerful framework to identify genetic determinants underlying complex plant–herbivore interactions, as demonstrated in Arabidopsis and other systems (Atwell *et al*., 2010; Kloth *et al*., 2016; Xu *et al*., 2023). Consistent with this, variation in aphid performance among accessions (**Fig. 1B**) was primarily associated with a locus on chromosome 2 identified by GWAS (**Fig. 2A; Supplementary Table S3**), encompassing members of the *CSLB* gene family (**Table 1**). Although *CSLB3* and *CSLB4* are in linkage disequilibrium (**Fig. 2B**), expression analyses following aphid infestation (**Fig. 3A**) support *CSLB4* as a likely contributor within this region. Additional loci on chromosome 4 also showed association signals (**Fig. 2A; Supplementary Table S3**), suggesting that aphid resistance is influenced by multiple genomic regions. While several CSL family members have been implicated in the biosynthesis of non-cellulosic polysaccharides (Dhugga *et al*., 2004; Farrokhi *et al*., 2006; Dwivany *et al*., 2009; Yang *et al*., 2020), the specific functions of the CSLB subfamily remain poorly characterized. Together, these findings identify *CSLB4* as a genetic component associated with plant–aphid interactions.

Although the effect size of the lead SNP appears modest (**Supplementary Table S3**), this is consistent with the quantitative nature of aphid resistance reported in Arabidopsis (Atwell *et al*., 2010; Kloth *et al*., 2016). In natural populations, allelic variation at individual loci often contributes incrementally to phenotypic variation, whereas functional perturbation in a controlled genetic background can reveal stronger phenotypic effects. Accordingly, loss-of-function mutants of *CSLB4* exhibited a clear reduction in aphid performance (**Fig. 4**), supporting the biological relevance of this locus. Together, these findings, along with the detection of additional associated loci, suggest that aphid resistance is shaped by multiple genetic components.

### CSLB4 modulates aphid performance and feeding behavior

Functional analyses of T-DNA insertion mutants demonstrated that disruption of *CSLB4* reduces aphid reproductive performance and alters host preference (**Fig. 4**), indicating reduced plant acceptance. Electrical penetration graph (EPG) analyses revealed that aphids reached phloem tissues more rapidly in mutant plants (**Table 2**), suggesting that *CSLB4* influences plant features encountered during early probing and phloem access phases. Despite earlier phloem access, aphid reproductive performance was reduced on *cslb4-1* plants. This observation indicates that enhanced resistance in *cslb4* mutants is unlikely to result from a physical barrier preventing stylet penetration, arguing against a purely mechanical constraint on aphid feeding. Instead, reduced aphid performance is more consistent with changes in feeding dynamics after vascular access is established, as supported by increased duration and frequency of salivation events (E1) and alterations in ingestion-related parameters (**Table 2**). Increased salivation is generally associated with difficulties in establishing sustained phloem ingestion (Tjallingii, 2006; Will and van Bel, 2006), suggesting less efficient feeding conditions and consistent with reduced phloem acceptance. Aphid-feeding behavior and performance on a given host are influenced by a combination of physical (Benatto *et al*., 2018), chemical (Kim *et al*., 2008), and nutritional traits (Nalam *et al*., 2021), including structural features such as tissue structure and organization, feeding site characteristics, as well as phloem composition (Powell *et al*., 2006; Escudero-Martinez *et al*., 2021). In this context, the observed changes in feeding behavior are consistent with altered host suitability rather than impaired access to vascular tissues. In addition, increased intracellular puncturing activity observed in mutant plants indicates altered tissue interactions during the exploratory phase (**Table 2**), consistent with modified stylet-tissue interactions that may influence feeding behavior. Together, these findings support a model in which *CSLB4* influences aphid performance through effects on feeding dynamics and host suitability, rather than through simple mechanical constraints on stylet progression.

Similar discrepancies between feeding behavior and reproductive success have been reported in other aphid systems, where insect performance depends not only on vascular access but also on host suitability following phloem ingestion (Lu *et al*., 2016; Nalam *et al*., 2021). Consistent with this, *CSLB4*-dependent plant traits influencing stylet-tissue interactions and feeding dynamics appear to affect not only stylet progression but also downstream processes that determine aphid success once phloem access is established. Taken together, these results support a decoupling between phloem accessibility and aphid performance, challenging the assumption that efficient phloem access directly predicts aphid performance.

### CSLB4 is associated with targeted remodeling of cell wall architecture

The conservation of glycosyltransferase-associated motifs (e.g., DXD, QXXRW, and TED) in CSLB4 (**Fig. 3B**) is consistent with a role in polysaccharide biosynthesis (Daras *et al*., 2021), although its precise enzymatic activity remains to be determined. Glycosyltransferases involved in XyG biosynthesis also relay on conserved catalytic features (Julian and Zabotina, 2022), suggesting functional parallels without implying a direct role for CSLB4 in XyG biosynthesis.

Biochemical analyses indicate that disruption of *CSLB4* leads to specific alterations in CW composition, particularly affecting pectic polysaccharides and monosaccharide ratios associated with rhamnogalacturonan I (RG-I) side chains (**Fig. 6A,C**), while no major changes in hemicellulose abundance were detected (**Fig. 6B**). These results suggest that loss of *CSLB4* does not cause large-scale changes in CW polysaccharide content but instead results in targeted changes in CW architecture. The observed shifts in pectic composition and RG-I–associated monosaccharide ratios are consistent with alterations in pectin architecture, a key determinant of CW organization and matrix interactions (Cosgrove, 2005; Atmodjo *et al*., 2013). RG-I side chains contribute to wall hydration, porosity, and the spatial arrangement of matrix polymers, and their remodeling has been associated with changes in cell adhesion and CW elasticity, likely through alterations in arabinan and interactions between pectins and cellulose (Arsovski *et al*., 2009). Pectins and xyloglucans (XyG) operate within an integrated CW matrix, where pectic domains can influence the accessibility and spatial arrangement of hemicelluloses relative to cellulose microfibrils (Park and Cosgrove, 2012a; Cosgrove, 2016). In this context, *CSLB4* disruption was associated with increased LM25 in mesophyll CWs (**Fig. 7**). Importantly, no differences in LM25 binding were detected in immunodot blot assays (**Supplementary Fig. S7A,B**), indicating that overall XyG epitope abundance is not altered. Together, these results are consistent with changes in XyG epitope accessibility and/or spatial distribution within the CW matrix rather than total XyG content, supporting a reorganization of polymer interactions within the CW matrix.

Although no direct measurements of mechanical properties were performed, previous studies show that changes in pectin structure, including side-chain composition and esterification status, can influence CW porosity and mechanics (Peaucelle *et al*., 2011; Gallemí *et al*., 2025), consistent with reduced pectin methylesterification and altered RG-I side-chain ratios in *cslb4-1* (**Supplementary Fig. S7C; Fig. 6C**;). Reduced pectin methylesterification can increase CW stiffness through Ca²□-mediated crosslinking of homogalacturonan (HG) (Wolf *et al*., 2009), potentially reinforcing the wall and limiting tissue penetration during pathogen attack (Lionetti *et al*., 2014). Together, our results indicate that CSLB4 primarily affects the spatial organization and interaction of matrix polysaccharides rather than their overall abundance. This interpretation is consistent with observations in other CW-related mutants and transgenic systems, in which targeted changes in matrix components such as XyG and RG-I can lead to pronounced biological effects (Park and Cosgrove, 2012b; Hassan *et al*., 2024). Notably, these architectural changes provide a plausible framework to explain the altered aphid-feeding behavior observed in *cslb4-1* (**Fig. 4**; **Table 2**), as local differences in CW organization are likely to influence the physical environment encountered during stylet progression.

### Cell wall remodeling influences plant responses to aphid feeding

In addition to mechanical effects, CW architecture is increasingly recognized as a determinant of plant interactions with pathogens and herbivores, acting not only as a structural barrier but also as a source of signals that activate defense responses (Hou *et al*., 2019; Wan *et al*., 2021). Structural changes in the CW can influence both resistance to penetration and the perception of CW–derived signals that trigger defense pathways (Bethke *et al*., 2016; Molina *et al*., 2021). In this context, the reduced callose deposition observed in *cslb4-1* following aphid infestation (**Fig. 8; Supplementary Fig. S9**) support a role for CSLB4 in modulating CW–associated defense responses. Notably, enhanced resistance occurs despite reduced callose accumulation, indicating that callose deposition is not directly associated with resistance in this plant-aphid system. This observation is consistent with the uncoupling between phloem access and aphid performance (**Table 2**; **Fig. 4C**) and suggests that additional CW-associated factors contribute to the phenotype.

Changes in CW organization may also affect plant responses by modulating the generation and perception of CW–derived signals and the activation of downstream defense pathways (Ellis *et al*., 2002; Hernández-Blanco *et al*., 2007; Bethke *et al*., 2016; Molina *et al*., 2021). CW remodeling has been proposed to engage CWI–related signaling mechanisms that regulate immune outputs (Hamann, 2015; Vaahtera *et al*., 2019). In line with this, pectin structure has been directly linked to plant defense against aphids, as HG methylesterification influences plant–aphid interactions (Silva-Sanzana *et al*., 2019). During early infestation, increased levels of de-methylesterified HG and calcium cross-linked pectic domains have been observed, highlighting the dynamic remodeling of pectin during aphid attack. Reduced pectin methylesterification may also facilitate the generation of oligogalacturonides (OGs), which act as damage-associated molecular patterns that activate defense responses (Ferrari *et al*., 2013; Bethke *et al*., 2014). Pectin-derived OG act as damage-associated molecular patterns that activate plant defense responses (Davidsson *et al*., 2017; Howlader *et al*., 2020), and exogenous OG application can induce salicylic acid–associated defense and callose deposition, enhancing resistance to aphids (Silva-Sanzana *et al*., 2022). Although the endogenous OG production during aphid-feeding remains to be demonstrated, these findings support a role for pectin-derived signals in modulating plant responses to aphids.

In *cslb4-1*, the combination of altered polysaccharide organization and modified aphid-induced responses is consistent with a shift in CW-associated signaling states, although direct evidence for altered CWI signaling remains to be established. While such mechanisms are well characterized in plant–pathogen interactions (Houston *et al*., 2016), their role in plant–insect interactions, particularly with phloem-feeding herbivores, is less understood. Within this framework, CSLB4-dependent CW remodeling may influence plant defense not by restricting aphid access but by shaping the physiological context of feeding, thereby affecting downstream defense responses that determine aphid performance.

### Implications for the role of cell wall architecture in plant–insect interactions

Together, our results indicate that CSLB4 likely contributes to plant susceptibility to the specialist aphid *B. brassicae* and highlight the importance of CW organization in modulating plant–insect interactions. Rather than causing large changes in CW polysaccharide abundance, CSLB4 appears to influence the spatial organization and epitope accessibility of CW components, which are likely to affect aphid -feeding behavior and plant defense responses. Notably, this effect is specific to *B. brassicae*, as no differences were observed in the performance of the generalist aphid *Myzus persicae*. This species-dependent response suggests that CSLB4-mediated CW remodeling does not confer broad-spectrum resistance but instead modulates plant–aphid interactions in a context-dependent manner. Specialist aphids, which are highly adapted to their host plants, may rely on finely tuned interactions with host tissues (Hogenhout and Bos, 2011), including CW architecture and phloem composition, and thus may be more sensitive to subtle structural changes. In contrast, generalist aphids may exhibit greater plasticity in accommodating variation in host traits (de Vos *et al*., 2007), potentially reducing sensitivity to CW perturbations. In this context, the induction of *CSLB4* upon aphid infestation, together with its association with increased susceptibility, raises the possibility that CSLB4-dependent CW remodeling is exploited by *B. brassicae* to facilitate feeding.

Overall, our results identify CSLB4 as a modulator of CW architecture that influences plant–aphid interactions by decoupling phloem accessibility from host suitability. These findings highlight the CW as a dynamic interface integrating structural and signaling functions and suggest that processes beyond initial stylet penetration contribute to aphid performance. Future work addressing the biochemical activity of CSLB4 and its role in hemicellulose-related processes will be essential to clarify the molecular mechanisms underlying this interaction.

## Conclusions

Our study identifies *CSLB4* as a regulator of plant–aphid interactions, linking cell wall (CW) organization to insect performance. By integrating GWAS with functional and physiological analyses, we show that loss of *CSLB4* enhances resistance to *Brevicoryne brassicae* while facilitating earlier access to the phloem, revealing a decoupling between phloem accessibility and host suitability. EPG analyses indicate that aphids reach vascular tissues more rapidly in *cslb4* mutants, but exhibit altered probing and salivation dynamics, suggesting that factors beyond initial stylet penetration contribute to feeding success. At the cellular level, *CSLB4* disruption leads to subtle but spatially distinct changes in CW organization, including altered xyloglucan epitope accessibility, modified pectin composition, and reduced aphid-induced callose deposition. These changes do not reflect major shifts in polysaccharide abundance but rather indicate targeted changes in CW architecture. Together, our findings identify *CSLB4* as a modulator of CW organization that influences plant susceptibility by affecting both early feeding behavior and downstream determinants of aphid performance. More broadly, this work highlights the CW as a dynamic interface integrating structural and physiological processes that shape plant–insect interactions.

## Supplementary Data

The following supplementary data are available at JXB online.

**Supplementary Table S1**. *Arabidopsis thaliana* accessions used in this study, including geographic origin, coordinates, and ABRC stock numbers.

**Supplementary Table S2**. Primer sequences used for RT–PCR analysis.

**Supplementary Table S3**. Top GWAS-associated SNPs for aphid performance

**Supplementary Table S4**. Haplotype composition of candidate genes identified in GWAS-associated genomic regions.

**Supplementary Figure S1**. Experimental design of the no-choice bioassay used for GWAS phenotyping.

**Supplementary Fig. S2**. Tissue-specific expression patterns of marker genes used for cell-type enrichment validation.

**Supplementary Fig. S3**. Population structure and phenotypic variation across the *Arabidopsis thaliana* panel.

**Supplementary Fig. S4**. Tissue-specific expression patterns of *CSLB3* and *CSLB4*.

**Supplementary Fig. S5**. Structural modeling of CSLB proteins.

**Supplementary Fig. S6**. Aphid performance of the generalist aphid *Myzus persicae* on *cslb4-1*.

**Supplementary Fig. S7**. Immunodot blot profiling and pectin methylesterification in cell wall fractions.

**Supplementary Fig. S8**. Additional confocal images of LM25 immunolabeling in leaf tissues.

**Supplementary Fig. S9.** Additional images of callose deposition following aphid infestation.

## Acknowledgements

We thank Sandy Rojas for assistance with sowing and plant maintenance. We are grateful to Adrián Moreno for providing the subcellular marker constructs. We also acknowledge the Centro de Biotecnología Vegetal for providing the facilities where this research was conducted.

## Author Contributions

FM and FB-H: conceptualization; FM, DA-G, DS, JD-R, IF-V, CI-A, MP-M, RV-S, DZ, and CF: methodology; FM: formal analysis; FM, DA-G, JD-R, CI-A, MP-M, MR, FO, and DS: investigation; FM, AH-V, SS-A, and FB-H: resources; FM: data curation; FM and FB-H: writing - original draft; FM, DS, MP-M, CF, AH-V, SS-A, and FB-H: writing-review & editing; FM, RV-S, and CF: visualization; FB-H: supervision; FM and FB-H: funding acquisition.

## Data Availability

The data of this study are available from the corresponding author upon reasonable request.

